# Standard treatment against paediatric BRAF-V600E glioma promotes senescence and sensitizes tumours to BCL-xL inhibition

**DOI:** 10.64898/2026.01.13.698644

**Authors:** Romain Guiho, Hiba Hamdi, Grace Cooksley, Timoty H. Chang, Lei Cao, Romain Sigaud, Ashley Vardon, Scott Haston, Florian Selt, Akang Bassey, Daniela Kocher, Daniel Gharai, Jesús Gil, Olaf Witt, Till Milde, Darren Hargrave, Pratiti Bandopadhayay, Paula Alexandre, Juan Pedro Martinez Barbera

**Affiliations:** Developmental Biology and Cancer Programme, Birth Defects Research Centre, Great Ormond Street Institute of Child Health, University College London, London, UK; Nantes Université, Oniris, INSERM, Regenerative Medicine and Skeleton, RMeS, UMR 1229, F-44000 Nantes, France; Department of Pediatric Oncology, Dana-Farber Cancer/Boston Children’s Cancer and Blood Disorders Center, Boston, MA, USA; Department of Neurosurgery, Beijing Tiantan Hospital, Capital Medical University, Beijing, China; University Hospital Jena, Department of Pediatrics and Adolescent Medicine, Friedrich Schiller University Jena, Jena, Germany; Comprehensive Cancer Center Central Germany (CCCG), Jena, Germany; Institute of Cancer and Genomic Medicine, University of Birmingham, Birmingham, UK; Clinical Trial Unit (ZIPO), Department of Pediatric Hematology, Oncology, Immunology and Pulmonology, Heidelberg University Hospital, Heidelberg, Germany; National Center for Tumor Diseases (NCT), Heidelberg, Germany; MRC Laboratory of Medical Sciences (LMS), Du Cane Road, London, W12 0NN, UK; Institute of Clinical Sciences (ICS), Faculty of Medicine, Imperial College London, Du Cane Road, London W12 0NN, UK; Hopp Children’s Cancer Center Heidelberg (KiTZ), Heidelberg, Germany; Clinical Cooperation Unit Pediatric Oncology, German Cancer Research Center Heidelberg (DKFZ), Heidelberg, Germany; Great Ormond Street Hospital NHS Trust, London, United Kingdom

**Author notes:** **Corresponding authors:** Romain Guiho –, Nantes Université, Oniris, INSERM, Regenerative Medicine and Skeleton, RMeS, UMR 1229, F-44000 Nantes, France., Prof. JP Martinez-Barbera, Developmental Biology and Cancer Programme, Birth Defects Research Centre, Great Ormond Street Institute of Child Health, University College London, London, UK. Equal contributors (Romain Guiho, Hiba Hamdi).

**Keywords:** BRAF-V600E driven glioma, BT-40, MAPK inhibitors, Navitoclax, senescence

## Abstract

**Background:** Chemo-, radio-, and targeted therapies can induce senescence in tumour cells, a process called therapy-induced senescence (TIS), which can be exploited therapeutically using compounds that kill senescent cells (senolytics). Children with BRAF-V600E activated brain tumours are commonly treated with Dabrafenib, Trametinib or Vinblastine (alone or in combination) with good responses. However, a subset of patients experiences tumour regrowth upon treatment withdrawal. Here, we explore the role of TIS and senolytic therapies in preclinical models of paediatric BRAF-V600E driven brain tumours.

**Methods:** Human BT-40 tumour cells derived from pleomorphic xanthoastrocytoma are used in vitro or in vivo orthotopically transplanted into mice and treated with either Dabrafenib, Trametinib or Vinblastine. Senescence is assessed by immunostaining and RNA sequencing. Sensitivity to senolytics is determined in vitro in dose response assays and in vivo.

**Results:** Trametinib treatment induces a senescent phenotype in BT-40 cells in vivo with activation of a senescence-associated secretory phenotype (SASP) and expansion of IBA-1 microglia. Senescence is induced in BT-40 cells in vitro when treated not only with Trametinib, but also with Dabrafenib and Vinblastine. In vitro testing identifies Navitoclax as a potent senolytic against BT-40 senescent cells, mostly through inhibition of the anti-apoptotic protein BCL-xL. In BT-40 tumour bearing mice, combination of Trametinib and Navitoclax causes the ablation of senescent BT-40 cells with significant reduction in tumour regrowth after treatment withdrawal and increased mouse survival.

**Conclusions:** This study shows that Navitoclax enhances the anti-tumour efficacy of Trametinib and limits tumour regrowth following treatment withdrawal.

**Key Points:**

- Trametinib, Dabrafenib and Vinblastine induce senescence in BT-40 cells in vitro
- Senescent BT-40 cells are sensitive to BCL-xL inhibition
- Combination of Trametinib and Navitoclax reduces rebound tumour growth in vivo

**Importance of the Study:** Children with BRAF-V600E activated glioma are frequently treated with Dabrafenib, Trametinib or Vinblastine with good responses. However, a subset of patients shows tumour regrowth when the treatment is stopped. To prevent this rebound growth, patients must continue treatment with potential aggravation in quality of life and risk of malignant progression. Our research demonstrates that these therapies induce BT-40 tumour cells to enter a non-proliferative state that shows features of cellular senescence. We identify that senescent tumour cells can be killed using Navitoclax, and potent inhibitor of the anti-apoptotic protein BCL-xL and a well-studied senolytic. We demonstrate that the combination of Trametinib and Navitoclax reduces tumour regrowth after treatment withdrawal. Our research provides a strong rationale supporting the combined use of senolytics with current conventional and targeted therapies against BRAF-V600E mutated brain tumours.

## Introduction

Brain tumours represent the most prevalent solid malignancies in children and the leading cause of cancer-related mortality in this population, with low-grade gliomas (LGG) accounting for 30–40% of central nervous system (CNS) tumours^1^. Clinical manifestations vary widely in LGG, primarily due to the compressive effects of the tumour on adjacent brain tissue or cerebrospinal fluid flow obstruction, often presenting with seizures, headaches, or alterations in cognition and behaviour. These tumours typically exhibit slow progression, limited invasiveness, and a low Ki-67 proliferation index^2,3^.

Genomic analyses have identified *KIAA1549::BRAF* fusions as the most frequent alteration in paediatric pilocytic astrocytomas, whereas BRAF-V600E mutations are common in more aggressive brain tumours such as diffuse astrocytomas, glioblastomas, pleomorphic xanthoastrocytomas, and gangliogliomas^4–9^. Treatment strategies depend on resectability and may involve surgery, chemotherapy (e.g. vinblastine), MAPK-pathway inhibitors (MAPKi)^10^ or radiotherapy. Initial preclinical studies using the BT-40 cell line, which was isolated from a patient with pleomorphic xanthoastrocytoma (PXA) harbouring the BRAF-V600E mutation and *CDKN2A/B* deletion, demonstrated the efficacy of MEK inhibitors against these tumours^11,12^. Subsequent clinical trials confirmed these findings, and recent investigations have further validated the use of combined BRAF and MEK inhibition, exemplified by dabrafenib and trametinib, against both fusion-positive and BRAF-mutant tumours, leading to regulatory approval for paediatric BRAF-V600E-driven gliomas^13–16^. Despite the good responses elicited by these MAPKi, a subset of patients shows tumour regrowth following discontinuation of treatment^17,18^. Although the reasons underlying this rebound growth remain unknown, rapid reactivation of the MAPK pathway has been observed in vitro, suggesting an intrinsic rebound mechanism within tumour cells. Additionally, elevated secretion of cytokines that recruit microglia may influence treatment responsiveness and post-therapy tumour regrowth^19^.

Senescence represents a cellular defence mechanism triggered by various stressors, including oncogenic signalling, replicative exhaustion, and DNA damage. Once senescence is initiated, cells undergo a stable proliferation arrest, orchestrated by key regulatory pathways such as p53/p21 and p16/RB. Beyond cell cycle exit, senescent cells exhibit non-cell-autonomous effects through the senescence-associated secretory phenotype (SASP). The SASP comprises a diverse array of secreted factors, including pro-inflammatory cytokines/chemokines and growth factors that can influence neighbouring cells and the tumour microenvironment^20,21^. Notably, conventional and targeted cancer therapies, such as radiotherapy, chemotherapy and molecular inhibitors, can promote therapy-induced senescence (TIS) in tumour cells. Residual senescent cells post-treatment may paradoxically facilitate tumour recurrence^22,23^. This can occur through cell-autonomous mechanisms, such as reactivation of proliferation due to p53/p21 or p16/RB pathway inactivation, or via paracrine signalling driven by persistent SASP activity^24,25^. Although oncogene-induced senescence has been investigated in LGG, the effects of current standard of care therapies in inducing senescence has not been thoroughly investigated ^19,26,27^.

We sought to investigate the role of TIS in contributing to tumour regrowth after treatment withdrawal using the well-validated BT-40 PXA tumour cells. Our data supports the notion that current treatments against paediatric BRAF-V600E-activated glioma tumour can induce senescence. We identify that a combination therapy of Trametinib plus Navitoclax is effective in reducing rebound growth after treatment withdrawal.

## Materials and Methods

### Cell Culture

BT-40 cells represent a uniquely relevant human model of paediatric low-grade glioma, carrying BRAF-V600E together with CDKN2A/B loss, a combination linked to aggressive behaviour and therapeutic vulnerability to MAPK pathway inhibition. Notably, BT-40 has been extensively used to identify and validate MAPK-targeted therapies now employed clinically. Given the scarcity of robust, genetically appropriate pLGG cell lines, BT-40 provides a clinically faithful and translationally meaningful platform for this study. BT-40 cells^12^ (kind gift of Dr. P. Houghton, University of Texas) were grown in RPMI1640 GlutaMAX (Gibco) supplemented with 10% foetal bovine serum (Gibco). BT-40 cells expressing LucF and GFP (referred to as BT-40-LucF) were generated by lentiviral strategy using pSLIEW vector with the previously described protocol ^28^. Extended protocol is available in Supplementary Material.

### Reagents

DMSO (Santa Cruz Biotechnology [SCB]) was used as vehicle control; Dabrafenib (MedChemExpress [MCE] - HY-14660), Trametinib (ApexBio - A3018), Vinblastine (SCB - sc-491749), Navitoclax (ABT-262; ApexBio - A3007), A-1155463 (MCE - HY-19725), A-1331852 (MCE - HY-19741), Obatoclax (MCE - HY-10969), Venetoclax (ABT-199; ApexBio - A8194), Piperlongumine (ApexBio - A4455), Digoxin (ApexBio - B7684). Characteristics of the BH3 mimetics used in this study are presented in **Supplementary Table S1**.

### Intracranial xenograft mouse model

Six-week-old female NOD scid gamma mice, purchased from Charles River UK, were used to establish intracranial xenografts. A total of 6 × 10^5^ cells BT-40 expressing LucF were injected into the right striatum of each mouse using a stereotactic frame (XYZ coordinate from bregma: −1.5mm; −1.5mm; 3+1mm). Extended protocol is available in **Supplementary Material**. Bioluminescence imaging (Firefly D-Luciferin, s.c 150mg/kg – PerkinElmer; IVIS Lumina III In Vivo Imaging System - PerkinElmer) was taken at day 12 after implantation and then every four days to monitor intracranial tumour growth. The mice were sacrificed when they exhibited weight loss >15% body weight or neurological symptoms. All mouse procedures were performed under licence, following UK Home Office Animals (Scientific Procedures) Act 1986, ARRIVE guidelines, and local institutional guidelines (UCL ethical review committee). A humane endpoint was chosen to avoid severe adverse effects and minimise suffering of mice harbouring tumours, according to UK Home Office and local Ethical Committee regulations. Brightfield and GFP of whole-mount brains were assessed with Leica M205 FA Fluorescence stereo microscope.

### FFPE Immunohistochemistry and Immunofluorescence

Immunostaining on 4µm formalin-fixed paraffin-embedded sections were performed as previously described^29^. At least three tumours and between five to six non-consecutive slides were analysed. As negative controls, sections were incubated with secondary antibody alone. A list of the antibodies and the antigen retrieval methods used is presented in **Supplementary Table S2**. Imaging was performed on Zeiss Observer 7 colour fluorescence, with Hamamatsu Flash 4.0v3 camera, using Zen Blue software v3.7. Quantification was performed with quPath software 0.5.1^30^.

### Cell viability, drugs combinatorial effects and apoptosis

Two thousand cells per well were seeded into 96-well plates and cultured for 72 h in the presence of increasing concentrations of drugs as indicated. Cell confluency was monitored by taking 3 brightfield pictures per well twice a week for 5 weeks. The BT-40 cells growing in large spindle-shaped clusters, confluency is represented as a ratio of measured covered cell area vs. maximum covered area. Cell viability was determined by CellTiter-Glo One Solution Assay (Promega) on a FLUOstar OPTIMA plate reader (BMG Labtech) 72h after treatment. IC50 were calculated using GraphPad Prism v9.4.0 software modelization. Drug sensitivity scores (DSS) were estimated using Breeze-pipeline^31^. Cell apoptosis was evaluated using Caspase-Glo 3/7 Assay (Promega), results were normalized by DAPI-positive nuclei counted prior to Navitoclax addition under the same treatments conditions. Extended protocol is available in **Supplementary Material**.

### Bulk RNA-sequencing analysis

For in vitro experiment, total RNA was extracted using RNeasy Mini kit (Qiagen) as previously described ^32^. Library preparation was performed with KAPA mRNA HyperPrep kit. Sequencing was then performed on an Illumina with NextSeq2000 100cycles P2 kit. Sequencing data was then aligned with {STAR 2.7.11b}. Raw and processed data are available on GEO repository GSE294664. Secondary analysis was performed using {R 4.3.3} on RStudio (2023.06.0+421). Differential expression was performed using {DESeq2 1.38.1}, complete pipeline is available at https://gitlab.univ-nantes.fr/guihomics/plgg_tis_rnaseq. GSEA performed using the GSEA software (v4.3.2 - UC San Diego and Broad Institute). Gene sets used in this study were either obtained from public databases: gene ontology (GO), KEGG, REACTOME, molecular signatures database (Broad Institute); or published literature (senescence profiling and SASP^33,34^).

### In vitro histochemistry and immunofluorescence

EdU incorporation staining was performed using Click-iT EdU Cell Proliferation Kit (Invitrogen) following manufacturer’s instructions. Senescence-associated (SA)-β-galactosidase staining was performed using the Senescence Cells Histochemical Staining Kit (CS0030 - Sigma-Aldrich) following manufacturer’s instructions. Immunostaining of adherent cells was performed after 10min fixation with Formalin 4% (v/v), permeabilization with 0.1% Triton X-100 and blocking with 2% BSA. Primary antibodies were incubated for 2 hours at RT before fluorescent dye–labelled secondary antibody incubation for 45min. DAPI was used as counterstain. The list of the antibodies used is presented in **Supplementary Table S2**. Imaging was performed on Zeiss Observer 7 colour fluorescence, with Hamamatsu Flash 4.0v3 camera, using Zen Blue software v3.7. Analysis was performed with Fiji software 1.53t^35^.

**Statistical Analysis**

Each experiment was repeated independently 3 times. Results are given as a mean ± SD for in vitro experiments and mean ± SEM for in vivo experiments. They were compared using unpaired t test or two-way ANOVA followed respectively by Bonferroni’s post-test or Tukey’s multiple comparisons test, calculated with GraphPad Prism v9.4.0 software. Kaplan-Meier results were computed using {ggsurvfit v1.1.0} R package. Results with p < 0.05 were considered significant and represented by an asterisk (*), p < 0.01 represented by two asterisks (**) and p < 0.001 represented by three asterisks (***).

## Results

### Trametinib induces cell senescence and activates a microglia attractant SASP in BT-40 tumour-bearing mice

First, we sought to test the effects of MAPK pathway inhibition in vivo using the clinically approved MEK inhibitor Trametinib. LucF-GFP BT-40 cells were orthotopically engrafted into NOD scid gamma (NSG) mice and tumour development was assessed by bioluminescence imaging (BLI). Once tumours were established (BLI> 3.10^6^ p.sec^−1^.cm^−2^.sr^−1^), mice were divided into two groups of equivalent BLI and treated with either Trametinib (1mg/kg) by oral gavage for 7 days (experimental group) or with vehicle (control group) (**Figure 1A**). As expected, Trametinib treatment led to a significant reduction in BLI compared with both the vehicle-treated group at the end of treatment (Day 14, Figure 1B; p=0.0009) and with the BLI at treatment initiation (Day 0, Figure 1B; p=0.0319). However, one week after treatment withdrawal, BLI levels in the Trametinib group increased again, indicating tumour regrowth (Figure 1B).

**Figure 1.**
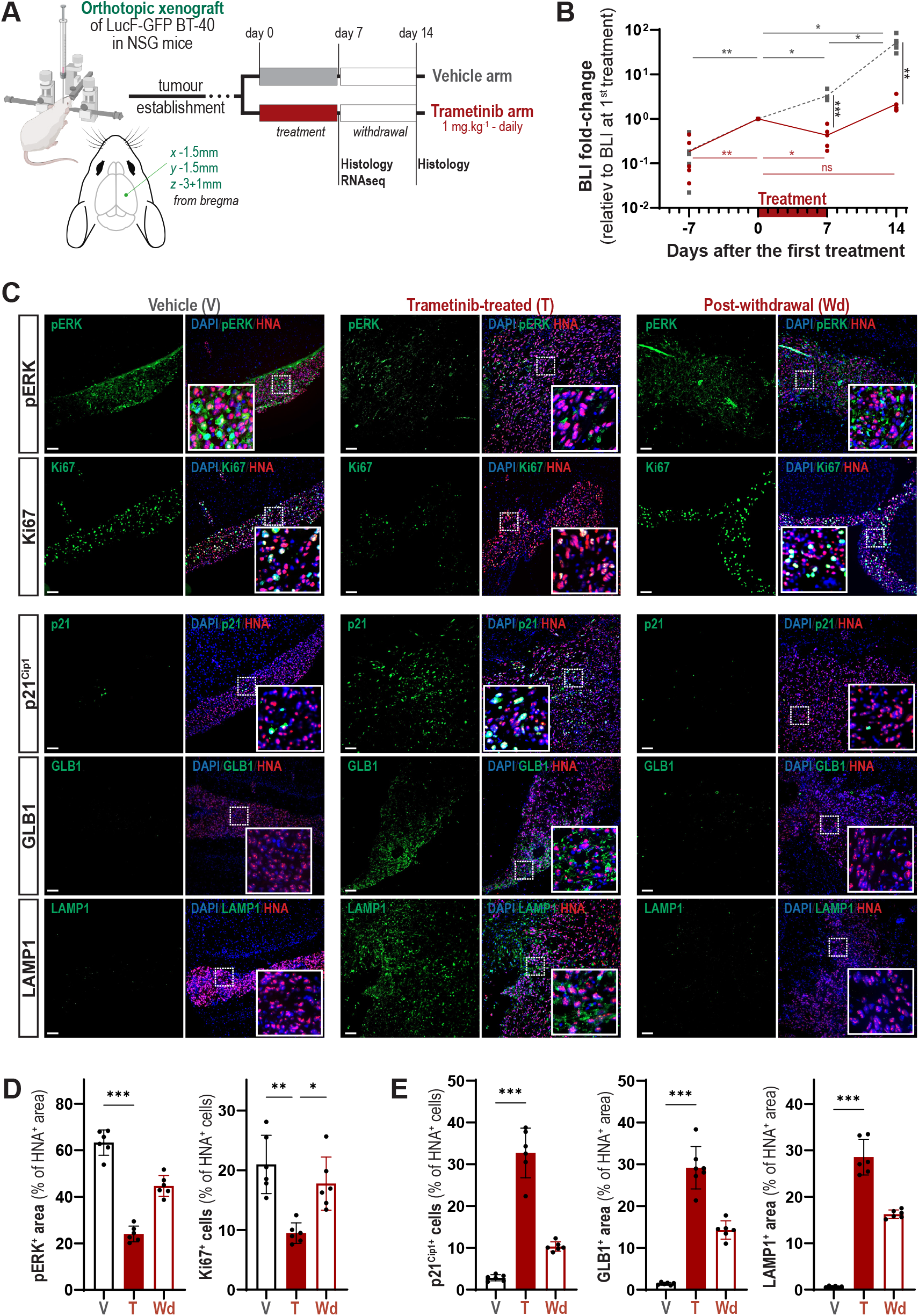
Trametinib treatment induces senescence and activates the SASP in BT-40 brain tumours in vivo. (A) A schematic of the experimental design. BT-40 tumour-bearing NSG mice are treated either with Trametinib or vehicle for 7 days, and analyses are performed at day 7 and day 14 (7 days post-Trametinib withdrawal). (B) Tumour growth is measured by bioluminescence imaging and is shown as mean values for 5 individual animals per group as radiance in photons.second^−1^.cm^−2^.steradian^−1^ relative to the radiance at day 1 of treatment. Note that although Trametinib treatments leads to a significant reduction in BLI at day 7, there is a quick rebound growth after treatment discontinuation at day 14. Two-way RM ANOVA Geisser-Greenhouse’s correction, Šídák’s multiple comparisons test between groups at indicated timepoints are indicated vertically, and differences between timepoints for Trametinib group are indicated horizontally. (C) Double immunofluorescence against Human Nuclear Antigen (HNA – red) and either phosphorylated-ERK1/2, Ki67, or the senescent markers p21, GLB1 or LAMP1 (green). DAPI in blue. Scale bars: 200 μm. (D,E) Quantification of the proportion of HNA positive cells or areas expressing the markers used in (C). Note the overall reduction of PERK1/2 and Ki67 (D) but overall increase in senescent marker expression upon Trametinib treatment (E). Data show mean ± SEM of n = 3 BT-40 tumours per marker. Kruskal-Wallis test, Dunn’s post-test; *: p<0.05, **: p<0.01, ***: p<0.001.

Next, we analysed BT-40 tumours by immunostaining for phosphorylated ERK1/2 (p-ERK1/2) as a readout of MAPK pathway activation, Ki67 as a proliferative marker, and HNA (human nuclear antigen) to identify BT-40 cells in mice treated with Trametinib for 7 days or examined 7 days after treatment withdrawal. Double immunofluorescence staining for p-ERK1/2 or Ki67 together with HNA revealed a significant reduction in p-ERK1/2-positive and proliferating BT-40 tumour cells in the Trametinib-treated group compared with vehicle controls (**Figure 1C, D**). As expected, following Trametinib withdrawal, the proportion of BT-40 cells expressing p-ERK1/2 and Ki67 increased markedly relative to the Trametinib-treated tumours, reaching statistical significance for Ki67 (**Figure 1C, D**).

To explore whether Trametinib treatment leads to senescence induction, we analysed the expression of: (i) the cell cycle inhibitor CDKN1A (p21^Cip1^), a key mediator of the senescence response (note that *CDKN2A*, encoding p16^INK4a^, another cell cycle regulator involved in senescence is deleted in BT-40 tumour cells); (ii) the lysosomal enzyme beta-galactosidase (encoded by the gene GLB1), responsible for the senescence-associated beta-galactosidase (SA-beta-gal) staining of senescent cells; and (iii) the lysosomal membrane protein LAMP1. The expression of these markers was negligible in the vehicle treated control tumours, but significantly elevated Trametinib treated tumours (**Figure 1C, E**). After drug withdrawal, senescent marker expression was noticeably reduced, which is consistent with the increased Ki67 staining previously described. These analyses suggest that Trametinib treatment results in MAPK pathway inhibition, reduction of proliferation and concomitant expression of senescent markers in BT-40 tumour cells, however, these responses are transient and diminish upon Trametinib withdrawal.

Next, we sought to validate this senescent phenotype using a molecular approach. BT-40 tumours were dissected from both Trametinib treated at day 7 and control vehicle groups, dissociated into single cell suspensions and tumour cells isolated by FACS and subjected to bulk RNA-sequencing. Although FACS-based isolation of BT-40 cells was limited by low GFP expression and therefore inefficient separation from murine brain cells, computational analysis of human RNA transcripts nevertheless enabled us to assess transcriptional variation within the samples. The data indicated substantial heterogeneity across BT-40 tumours, as reflected by the wide dispersion in the PCA plot (**Supplementary S1A**), with a discernible separation between vehicle- and trametinib-treated groups. Differential expression analysis reaffirmed previously reported trametinib-induced changes, including upregulation of CDKN1A and SASP-related genes and repression of proliferation-associated genes (**Supplementary S1B; Supplementary Table S3**). Geneset enrichment analysis (GSEA) revealed an enrichment for active MAPK pathway associated gene sets in vehicle control group highlighting an inhibitory effect of trametinib on the MAPK pathway in BT-40 treated tumours (**Supplementary Figure S1C**). Similarly, proliferation-associated gene sets (e.g. E2F targets and cell cycle) were found to be enriched in vehicle control tumours (**Supplementary Figure S1D)**. Of note, the SenMAYO geneset^34^, which contains a curated list of genes that characterises cellular senescence, as well as the SASP geneset^33^, were enriched in Trametinib-treated BT-40 tumours, supporting the hypothesis that MAPK pathway inhibition by Trametinib results in senescence induction in BT-40 cells in vivo (**Supplementary Figure S1E**).

Because the SASP can strongly influence immune cell recruitment, we examined SASP components in Trametinib-treated BT-40 cells. Among the factors upregulated, transcripts encoding the chemokines CCL2, CXCL10, and the cytokine IL6 were markedly increased in senescent BT-40 cells (**Supplementary Figure S1F**). Consistent with these transcriptomic data, double immunofluorescence for CCL2, CXCL10, or IL6 together with HNA showed a significant rise in the number of BT-40 tumour cells expressing each of these SASP cytokines/chemokines in Trametinib-treated mice compared with vehicle controls (**Figure 2A, B**). Following Trametinib withdrawal, expression levels trended downward but did not reach statistical significance (**Figure 2A, B)**.

**Figure 2.**
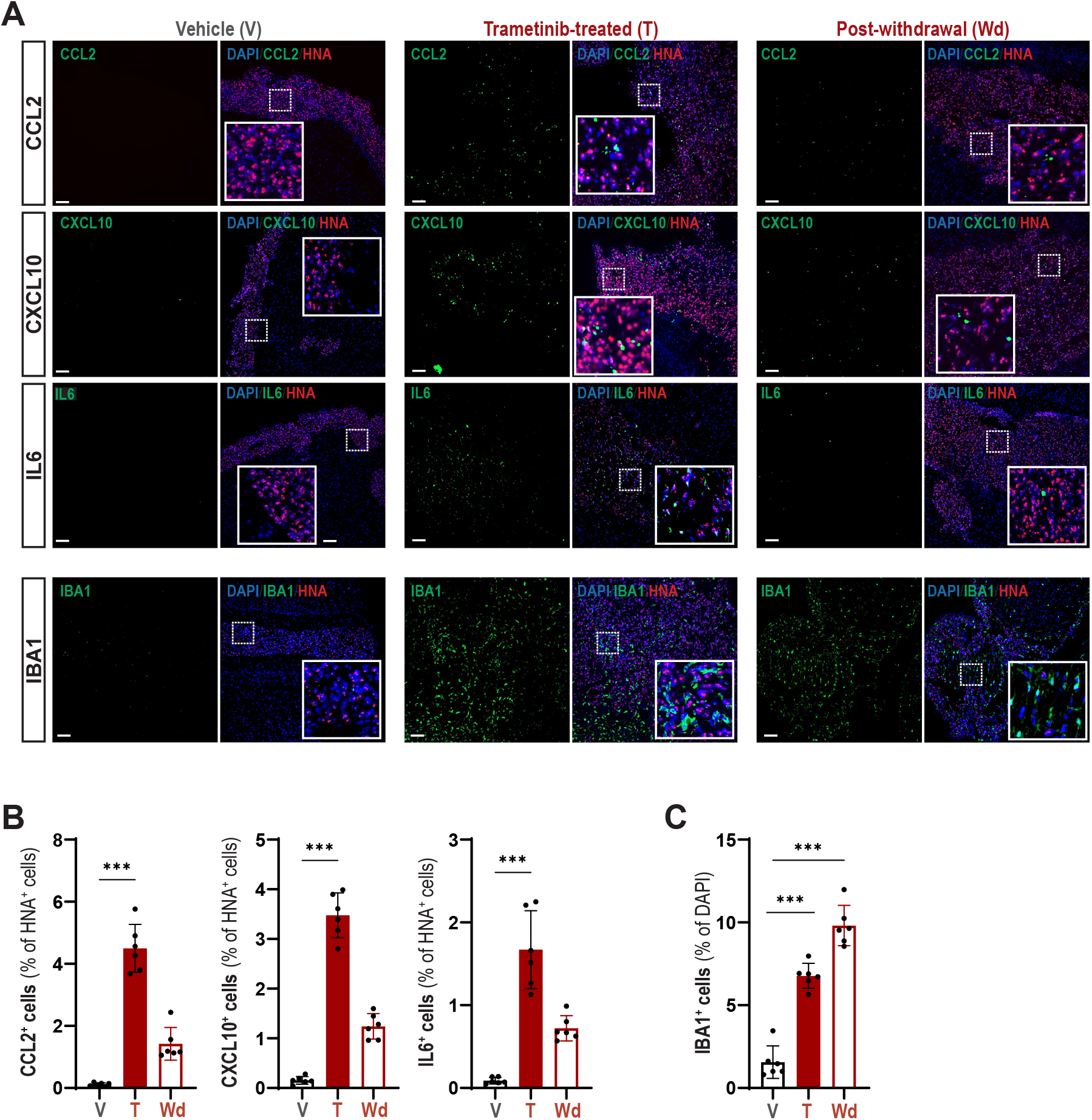
Trametinib-induced senescent BT-40 tumours express myeloid-attractant chemokines and cytokines, accompanied by prominent microglial infiltration. (A) Double immunofluorescence against Human Nuclear Antigen (HNA – red), the myeloid-attractant chemokines/cytokines CCL2, CXCL10 and IL6 (green) and the microglial/monocyte marker IBA1 (green). DAPI in blue. Scale bars: 200 μm. (B) Quantification of the proportion of HNA positive cells expressing the analysed markers in (A). Note the significant increase in marker expression upon Trametinib treatment. (C) Quantification of the proportion of cells expressing the IBA1 marker from (A). Note the significant increase in IBA1 positive cells expression upon and after Trametinib treatment. Data show mean ± SEM of n = 3 BT-40 tumours per marker. Kruskal-Wallis test, Dunn’s post-test; ***: p<0.001.

Given that CCL2, CXCL10, and IL6 play key roles in attracting and activating myeloid lineage cells^19^, we next assessed microglial infiltration. Immunofluorescence for IBA1 revealed abundant microglia in Trametinib-treated tumours, whereas vehicle-treated tumours contained only negligible numbers of IBA1-positive cells (**Figure 2A**). Notably, microglial abundance increased even further after Trametinib withdrawal, despite the accompanying reduction in SASP cytokine/chemokine expression (**Figure 2A, C)**.

Taken together, these findings indicate that Trametinib treatment not only suppresses proliferation and induces a senescence-associated programme, but also activates a SASP that expands the microglial compartment in BT-40 tumour-bearing mice.

### Trametinib, Dabrafenib, and Vinblastine are capable of inducing cellular senescence and activating the SASP in BT-40 cells in vitro

Next, we sought to compare how BT-40 cells respond in vitro to treatments commonly used for paediatric glioma, namely Trametinib, Dabrafenib, and Vinblastine. First, we carried out viability and apoptosis assays to identify the optimal conditions capable of inducing cell cycle arrest with minimal effects on cell viability and apoptosis. BT-40 cells were cultured at increasing drug concentrations ranging from 10^−10^ to 10^−6^ M, and apoptosis and cell viability determined at day 1 and 3 post-drug treatment, respectively (**Supplementary Figure S2A**). These studies revealed that drug concentrations up to 10^−7^ M (100 nM) caused minimal cell death or loss of cell viability (**Supplementary Figure S2B**).

We next examined how these drugs influence BT-40 cell proliferation using two complementary experimental designs. In an extended-treatment paradigm, cells were continuously exposed to Dabrafenib, Trametinib, or Vinblastine at 1 nM, 5 nM, or 10 nM for 7 days before drug withdrawal. In a second, short-term exposure paradigm, cells were treated with 100 nM of each drug for 16 hours, after which the drug was removed and cultures were maintained in drug-free media (Supplementary Figure S2C). We reasoned that these two paradigms model distinct stages of treatment, initial response during exposure and behaviour after withdrawal. In both settings, EdU incorporation was measured on day 7 as a readout of proliferation, and cell confluence was monitored until day 35.

Across both designs, all three drugs reduced EdU incorporation at 10 nM and 100 nM, consistent with cell-cycle arrest, whereas only Trametinib and Vinblastine reduced proliferation at 5 nM (Supplementary Figure S2D). After drug withdrawal, the durability of the arrest differed markedly by drug and concentration. Removal of Dabrafenib or Trametinib allowed BT-40 cells to resume proliferation within two weeks at all concentrations tested (5 nM, 10 nM, and 100 nM). By contrast, withdrawal of Vinblastine induced a much more stable arrest that persisted through day 35, when the experiment concluded (Supplementary Figure S2E).

Next, we characterized the senescent phenotype induced by these drugs molecularly and by immunostaining. Based on the previously described findings, we proceeded with the 16 hours treatment and wash-out paradigm for subsequent experiments. This design was selected because brief exposure (16 hours) to the drugs, followed by withdrawal, produced a more durable and homogeneous growth arrest, consistent with a stable senescence-like state. At day 7, SA-beta-gal and immunofluorescence staining revealed a significant increase of both SA-beta-gal and p21^CIP1^-expressing cells suggesting senescence induction (**Figure 3B,C**). To confirm this phenotype, we performed bulk RNA sequencing of BT-40 cells 7 days after short-term treatment with Dabrafenib, Trametinib or Vinblastine and vehicle control. As expected, the transcriptomic profiles of the treated cells were highly altered compare to the cells treated with the vehicle, as reflected by: (i) volcano plots (**Figure 3D**); (ii) separation of experimental groups in the PCA plots (**Supplementary Figure S3A**); and (iii) the unsupervised ranking of the 500 most differentially expressed genes among the different experimental groups (**Supplementary Figure S3B**). Gene pathway analysis revealed the enrichment of pathways associated with cell proliferation in the vehicle treated BT-40 cells (**Figure 3E**). In contrast, treatment with either of the drugs resulted in the significant enrichment of pathways related to senescence, SASP and SASP-associated biological processes (**Figure 3E**). Supporting the RNAseq data, immunofluorescence staining confirms the expression of IL1β in Dabrafenib and Vinblastine-treated cells, as well as CCL2 in Dabrafenib, Trametinib, and to a lesser extend Vinblastine-treated cells (**Figure 3F**). These observations are consistent with selected SASP component expression represented in heatmap (**Figure 3G**).

**Figure 3.**
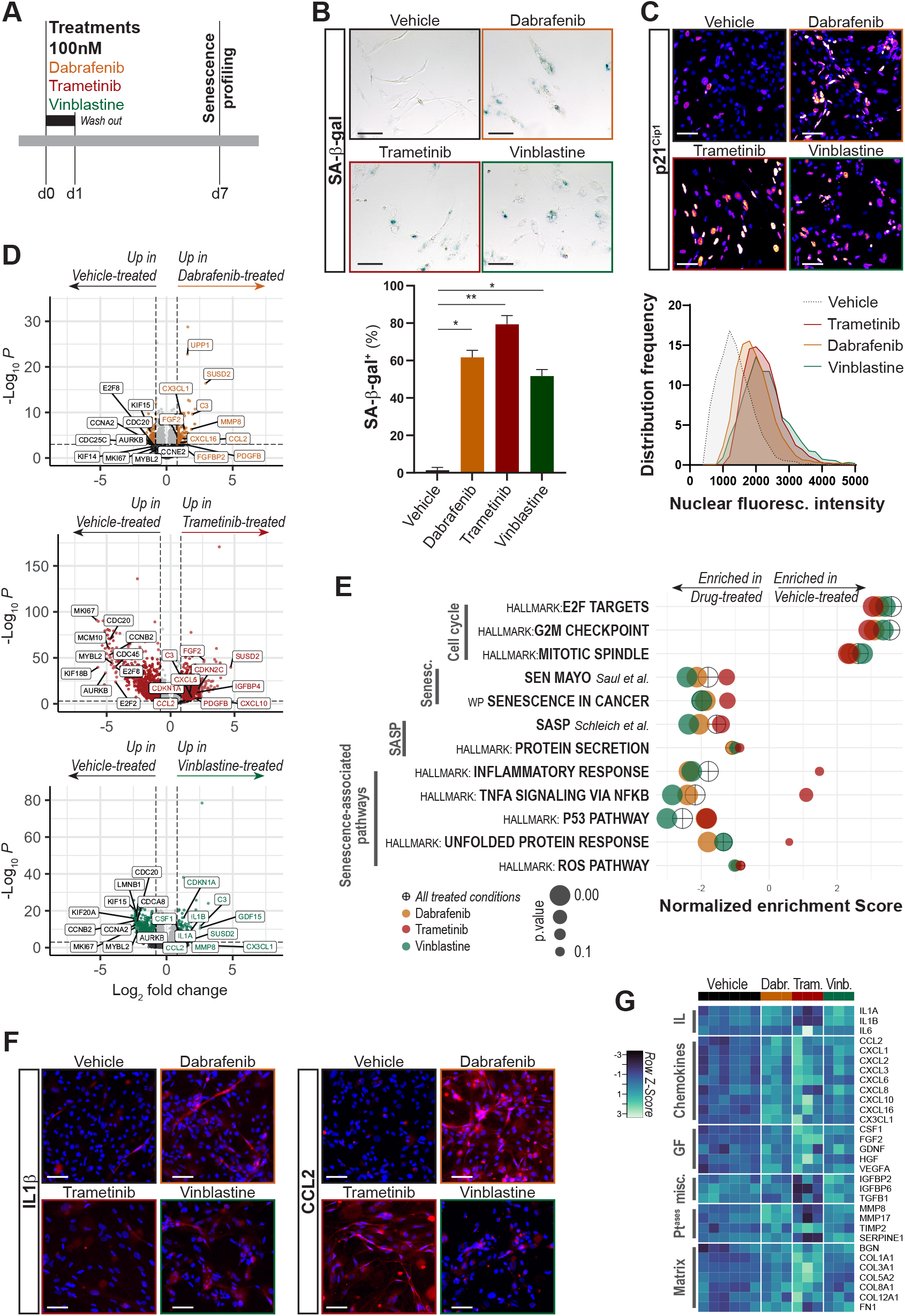
Treatment of BT-40 cells in vitro with Dabrafenib, Trametinib, or Vinblastine induces cellular senescence and activates the SASP. (A) Diagram of the experimental approach. Cultured BT-40 cells are treated with either of the drugs at day 0 for 24h, followed by a wash-out period of 6 days in the absence of drug, and analysed at day 7 both molecularly (RNA-seq) and by immunocytochemistry. (B) SA-β-Gal staining indicating the increase of blue-stained cells after drug treatment compared with vehicle-treated control. Kruskal-Wallis test, Dunn’s post-test; *: p<0.05, **: p<0.01. (C) p21^Cip1^ immunostaining showing the accumulation of nuclear fluorescence signal in the drug-treated cells compare with vehicle controls. (D) Volcano plot of differentially expressed genes comparing proliferative and drug-treated BT-40 cells. Thresholds used: p-values < 0.05 and log2 FoldChange > 1. (E) Normalized enrichment scores (NES) from GSEA showing decreased proliferation-associated genesets and increased senescence and SASP-related genesets for each of the treatment (Dabrafenib, Trametinib and Vinblastine) compared with vehicle control cells. The crossed dots represent the NES integrating the three treated conditions versus the vehicle control. (F) Immunofluorescence staining of IL1B and CCL2 (red) at day 7 in vehicle-treated and drug-treated BT-40 cultured cells. DAPI in blue. Scale bars: 100 μm. (G) Heat maps showing the relative expression of selected SASP factors in vehicle-treated and drug-treated BT-40 cells. Note that there are specific differences between treatments. IL: Interleukin, GF: Growth factors, Pt^ases^: matrix proteases. n=6 biological replicates for vehicle-treated BT-40, n=3 biological replicates for dabrafenib, trametinib and vinblastine-treated BT-40.

Although the three treatments were able to induce senescence and SASP, we noticed that there were differences in the dysregulated genes among treatments, suggesting drug-specific responses (**Figure 3D, Supplementary Figure S3C**). When comparing the transcriptomes of the differentially expressed genes (DEGs) between treated cells and controls, we identified a core signature comprising 132 genes shared between the three treatments, which included senescence and SASP genes such as *CDKN1A* (encoding the cell cycle inhibitor p21^CIP1^ - **Supplementary Figure S3D**), *SUSD2* (involved in negative regulation of cell cycle), and secreted factors such as *IL6, CCL2, PDGFB* and *VEGFA*. Trametinib-treatment induced drastic transcriptomic changes with 1363 specific DEGs, which included many lysosome-associated genes, suggesting lysosomal expansion, a hallmark of senescence (**Supplementary S3C and E**). Additionally, *CXCL6, CSF1, TIMP2* and matrix components *COL1A1, COL3A1* and *COL5A2* were specifically upregulated in Trametinib-treated cells (**Figure 3G, Supplementary Tables S4-S5-S6**). Other SASP components, e.g. *IL1A, IL1B, TGFB1* and metalloproteases *MMP8* and *MMP17* were upregulated in Dabrafenib and Vinblastine-treated cells, but not in Trametinib-induced senescence state. Together, these analyses demonstrate that while BT-40 cells treated with Trametinib, Dabrafenib, or Vinblastine display senescent signatures in vitro, the extent and pattern of SASP gene upregulation differ depending on the specific drug used.

### Senescent BT-40 tumour cells are highly sensitive to Bcl-xL inhibition

Senescence cells can often show specific vulnerabilities that may not be present in the same cells when in a non-senescent state. Therefore, we decided to explore whether the treatment with Trametinib, Dabrafenib or Vinblastine and induction of a senescent state in BT-40 cells could be exploited therapeutically using senolytics. We tested three described senolytic agents, Piperlongumine, Digoxin, and Navitoclax on senescence-induced (i.e. previously treated with either Dabrafenib, Trametinib or Vinblastine) and control proliferative, vehicle-treated BT-40 cells (**Figure 4A**). From these, only Navitoclax demonstrated a selective ablation of senescent cells in a dose response manner (**Figure 4B, C, Suppl. Figure S4**). Compared with proliferative BT-40 cells, senescent cells induced by Dabrafenib, Trametinib, or Vinblastine were 6-, 50-, and 7-fold more sensitive to Navitoclax, respectively (**Figure 4C**). Confirming these differences in responses to Navitoclax treatment, ssGSEA (single-sample GSEA) was used to calculate enrichment scores for each individual sample ^26,36^. ssGSEA revealed that treatment with either of the drugs resulted in higher Navitoclax-sensitivity and a lower Navitoclax-resistance relative to proliferative control BT40 cells (**Figure 4D**). Of note, the highest Navitoclax-sensitivity and lowest Navitoclax-resistance scores were observed in Trametinib-induced senescent BT-40 cells (**Figure 4D**).

**Figure 4.**
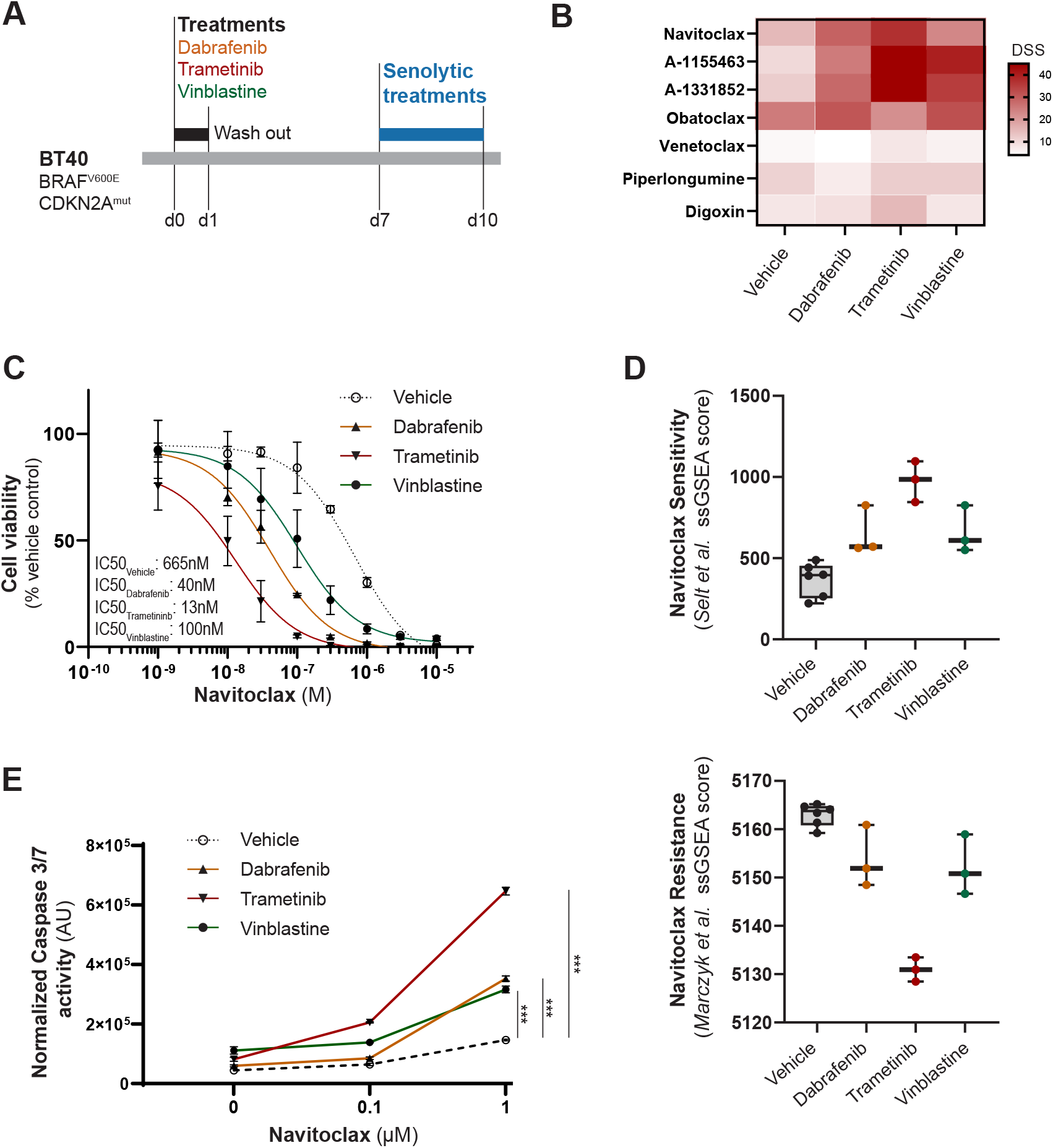
Treatment-induced senescent BT-40 cells are selectively vulnerable to Bcl-xL inhibition. (A) Schematic of the experimental design. Drug treatment is carried out for one day, followed by a wash-out period of 6 days in the absence of drug. At day 7, control proliferative and treatment-induced BT-40 cells are cultured in the presence of senolytics for 3 days. (B) Heatmap showing the drug sensitivity score (DSS) of several inhibitors of the Bcl-2 family of anti-apoptotic proteins and known senolytics used on cells pre-treated with vehicle, Dabrafenib, Trametinib or Vinblastine. Note that the highest DDSs are observed with Navitoclax, A-1155463, A-1331852 and Obatoclax, which all inhibit Bcl-xL, while the weak Bcl-xL inhibitor, but strong Bcl-2 inhibitor Venetoclax or the senolytics (Piperlongumine and Digoxin) show lower DSSs. (C) Dose response curves of Navitoclax on BT-40 cells pre-treated with either Dabrafenib, Trametinib, Vinblastine or vehicle control. Note that the IC50 values are lower in treatment-induced senescent cells than in proliferative BT-40 cells. (D) Navitoclax sensitivity (top: Selt_BT308.UP) and resistance (Marczyk_Navitoclax resistance) scores inferred from ssGSEA analysis of RNA sequencing data of drug-treated and vehicle controls. (E) Graph showing Caspase 3/7 enzymatic activity of treatment-induced senescent or proliferative, vehicle control BT-40 cells and cultured in the presence of 0.1uM or 1mM Navitoclax as shown in (A). Note the significant increase in senescent cells relative to proliferative control cells. AU Normalized activity by cell numbers. Kruskal-Wallis test, Dunn’s post-test; ***: p<0.001.

Navitoclax can inhibit Bcl-2, Bcl-xL, and Bcl-w ^37^. To identify the specific protein underlying the senolytic effect of Navitoclax, we tested chemical BH3 mimetics with specific inhibitory profiles. Drug sensitivity scores (DSS) were established from dose-response curves data obtained from drug-treated and control BT-40 cells and depicted in a heatmap (**Figure 4B**). Regardless of the specific drug used, we observed DSS lower than 15 for Venetoclax, a potent Bcl-2, but weak Bcl-xL inhibitor. In contrast, DSS values obtained for inhibitors with strong affinity to Bcl-xL such as Navitoclax, the pan-Bcl2 inhibitor Obatoclax or more relevant, the selective Bcl-xL inhibitors A-1155463 and A-1331852 were greater than 25, suggesting a higher sensitivity of treatment-induced senescent BT-40 cells to Bcl-xL inhibition (**Figure 4B, Supplementary Figure S4A-F**). Finally, we compared the IC50 values of Navitoclax in senescent BT-40 cells from our study with those reported in the Genomics of Drug Sensitivity in Cancer (GDSC) database, which encompasses 928 cell lines. Senescent BT-40 cells induced by treatment ranked among the most sensitive lines, with trametinib-induced senescence 2nd, dabrafenib-induced senescence 17th, and vinblastine-induced senescence 50th (**Supplementary Figure S4G**).

Finally, inhibition of BCL-xL is expected to result in apoptosis. To assess whether Navitoclax induces apoptosis in BT-40 senescent cells, senescent and proliferative cells were treated with Navitoclax (0.1µM and 1.0μM) or vehicle, and caspase 3/7 activity quantified. Treatment with Navitoclax resulted in increased effector caspase 3/7 activity, suggesting induction of apoptosis (**Figure 4E**). Together, these studies demonstrate that treatment-induced senescent BT-40 tumour cells are selectively vulnerable to apoptosis upon Bcl-xL inhibition, with Trametinib-treated cells showing the greatest sensitivity to Navitoclax.

### Combined Trametinib and Navitoclax therapy reduces tumour burden and prolongs survival in BT-40 tumour-bearing mice

We next evaluated the efficacy of combined Trametinib and Navitoclax therapy in BT-40 tumour-bearing mice. NSG mice were orthotopically transplanted with LUCF-GFP BT-40 tumour cells, and tumour growth was monitored by bioluminescence imaging (BLI). Once BLI reached 3.10^6^ p.sec^−1^.cm^−2^.sr^−1^, mice were evenly distributed into four groups with comparable tumour burdens: (1) Vehicle control, receiving the oral vehicle used for Trametinib and Navitoclax (n = 5); (2) Navitoclax, 50 mg/kg/day via oral gavage, 5 days per week for 2 weeks (n = 5); (3) Trametinib, 1 mg/kg/day via oral gavage, 5 days per week for 3 weeks (n = 7); (4) Combination therapy, Trametinib alone for 1 week, followed by Trametinib plus Navitoclax for 2 weeks, administered orally at the same doses and schedules (n = 9) (**Figure 5A**). All treatments were well tolerated, with no significant weight loss observed (**Supplementary Figure S5A**).

**Figure 5.**
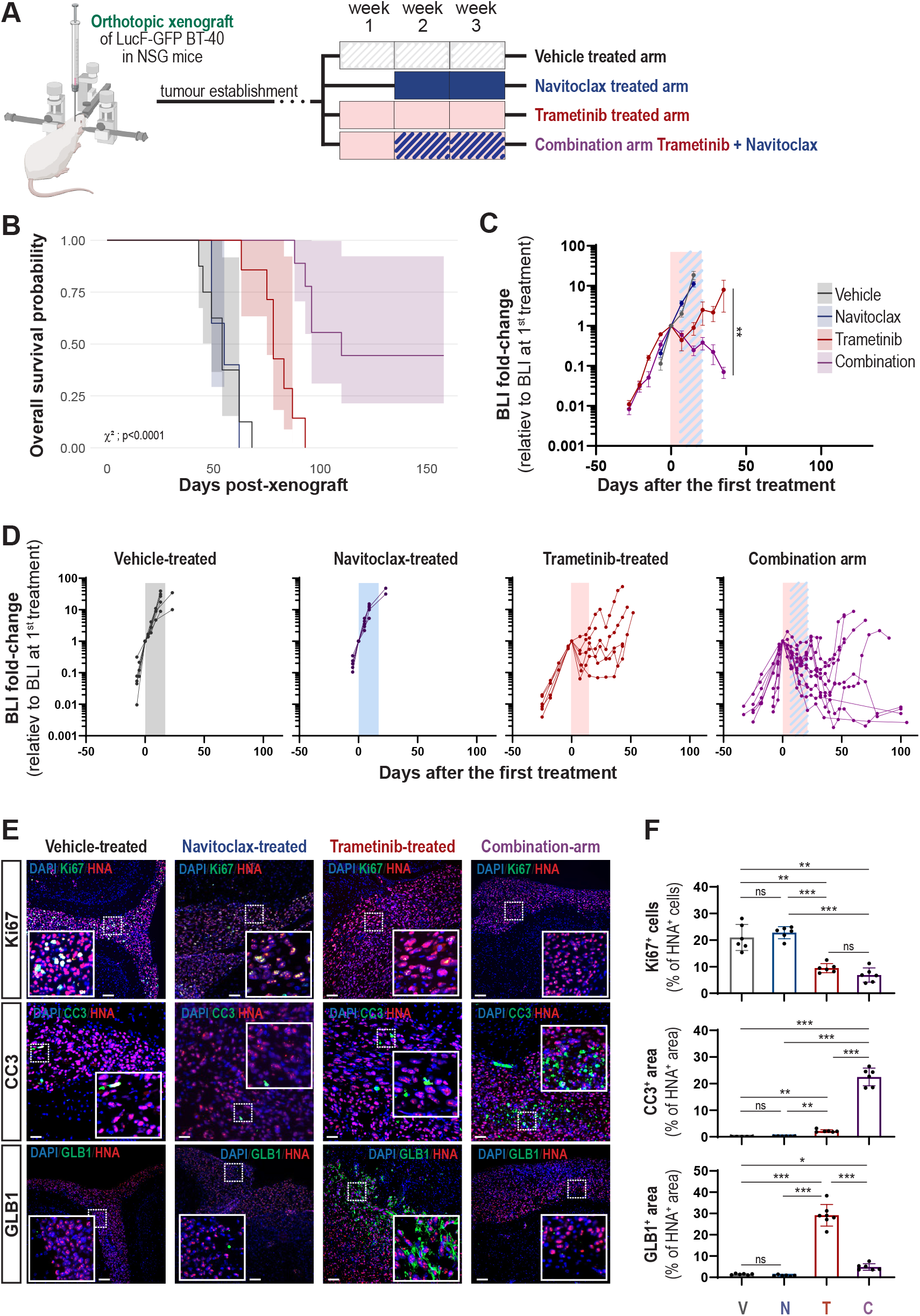
Combination therapy with Trametinib and Navitoclax demonstrates significant anti-tumour activity and extends survival in BT-40 tumour-bearing mice. (A) Schematic of the preclinical experimental design. Upon engraftment of BT-40 cells, tumour-bearing mice were split into 4 experimental groups: (i) vehicle treated for 3 weeks; ii) Navitoclax treated for 2 weeks (weeks 2 and 3); (iii) Trametinib treated for 3 weeks; (iv) Combination group in which mice were treated with Trametinib for 3 weeks and Navitoclax on weeks 2 and 3. (B) Kaplan-Meier estimator plots showing the probability of survival of the mice in the 4 different groups. Pairwise comparisons using Log-Rank test: Combi. vs. Veh.: p=0.0001; Combi. vs. Nav.: p= 0.0005; Combi. vs. Tram.: p= 0.0008; Tram. vs. Nav.: p= 0.0028; Tram. vs. Veh.: p= 0.0020. (C) Average BLI fold-change in the 4 experimental groups relative to the average BLI at day 1 of treatment. Tukey’s multiple comparisons test at day 15: Tram. vs. Nav.&Veh.: p <0.0001; Combi. vs. Nav.&Veh.: p <0.0001. Tukey’s multiple comparisons test at day 35: Tram. vs. Combi.: p=0.0028. (D) BLI fold-change for each individual mice in the 4 experimental groups relative to the BLI at day 1 of treatment. (E) Double immunofluorescence against Human Nuclear Antigen (HNA - red) and the proliferation marker Ki67, the apoptosis marker cleaved-caspase-3 (CC3), or the senescent marker GLB1 (green). DAPI in blue. Scale bars: 200 μm. (F) Quantification of the proportion of HNA positive cells expressing Ki67 and the percentage of tumour area positive for CC3 and GLB1 analysed in (E). Data show mean ± SEM of n = 3 BT-40 tumours per marker. Kruskal-Wallis test, Dunn’s post-test; *: p<0.05, **: p<0.01, ***: p<0.001.

Kaplan-Meier survival curves revealed a median survival of 54 days for the vehicle control group, 55 days for the Navitoclax-treated group and 78 days for the Trametinib group (**Figure 5B**). Remarkably, in the combination group, 4/9 mice survive until day 150 without showing any neurological symptoms. Pairwise comparisons using the log-rank test demonstrated significant differences in survival across treatment groups (**Figure 5B**). Representation of the average BLI fold change in the Trametinib and combination groups showed a significant reduction in BLI values (**Figure 5C**). Similarly, when BLI fold change was examined for each individual mouse in the preclinical study, the combination group displayed a variable treatment response: 5 of 9 mice showed an increase in BLI following treatment withdrawal, whereas 4 of 9 exhibited a remarkable response, with BLI values progressively decreasing to nearly undetectable levels (**Figure 5D, Supplementary Figure S5B**). These findings demonstrate significant anti-tumour activity of the Trametinib and Navitoclax combination against BT-40 tumours, albeit with intra-group heterogeneity in the duration of treatment efficacy.

We aimed to investigate the cellular and molecular mechanisms underlying the efficacy of the Trametinib plus Navitoclax combination. Tumours from the four experimental groups were analysed during treatment, using double immunofluorescence staining for proliferation, apoptosis, and senescence markers, with an anti-HNA antibody to specifically identify BT-40 tumour cells. Both, Trametinib only and the combination treatments resulted in a significant reduction in the proliferative index when compared with the vehicle control and Navitoclax-treated groups, as expected (**Figure 5E,F**). Conversely, cleaved caspase 3 expression, a marker of apoptosis, was significantly increased in the combination group relative to all other three groups (**Figure 5E**). Number of cells expressing senescent markers, such as p21^CIP1^, GLB1 and LAMP1 were all elevated in the Trametinib only group relative to the control group as previously shown (**Figure 1C,E, Figure 5E,F, Supplementary Figure S6A,B**). However, in the combination group senescent marker expression was significantly reduced compared with the Trametinib only group (**Figure 5E,F, Supplementary Figure S6A,B**).

Finally, we analysed the microglia compartment, as we have previously shown the presence of abundant IBA1+ microglia after Trametinib treatment. Double immunostaining against the IBA1 and HNA confirmed the presence of abundant IBA1-expressing microglia in the Trametinib-only treated group relative to both the vehicle control and Navitoclax-only groups (**Figure 6A,B**). Remarkably, microglia numbers were vastly reduced in the combination group relative to the Trametinib only group (**Figure 6C)**. The significant reduction in microglia numbers in the combination group was associated with a significant decrease in numbers of CCL2, CXCL10 and IL6-expressing cells (**Supplementary Figure S6C,D**). Together, these analyses align with our previous in vitro observations and confirm that Trametinib treatment induces senescence in BT-40 cells. Furthermore, our data indicate that Navitoclax can ablate senescent BT-40 tumour cells, resulting in decreased chemokine expression and reduced microglial infiltration into the tumours

**Figure 6.**
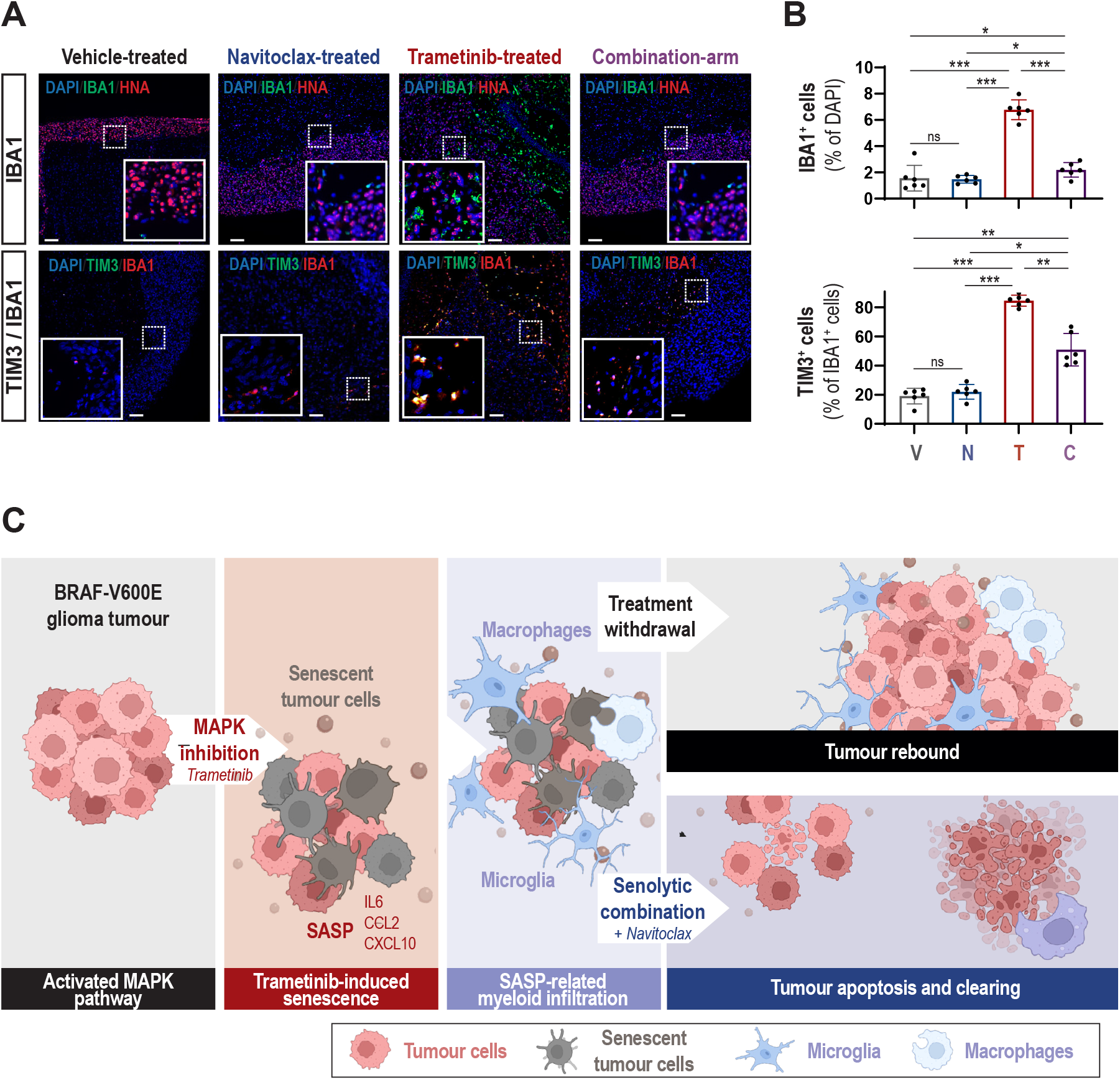
Combination therapy with Trametinib and Navitoclax reduces microglial infiltration in BT-40 tumours in vivo. (A)Double immunofluorescence against Human Nuclear Antigen (HNA – red) and microglial/monocyte markers (IBA1 and TIM3 - green). DAPI in blue. Scale bars: 200 μm. (B) Quantification of the proportion of DAPI positive cells expressing the stated markers from (A). Kruskal-Wallis test, Dunn’s post-test; *: p<0.05, **: p<0.01, ***: p<0.001. (C) Schematic model summarizing the responses of BT-40 cells to Trametinib or Trametinib plus Navitoclax treatment in BT-40 tumour-bearing mice. Trametinib induces robust senescence and activates the SASP, including myeloid-attractant factors (e.g., IL6, CCL2, CXCL10). Upon Trametinib withdrawal, BT-40 cells resume proliferation and myeloid infiltration increases. In contrast, combination therapy with Trametinib and Navitoclax selectively ablates senescent BT-40 cells, modulates the SASP, reduces myeloid infiltration, and promotes tumour clearance.

## Discussion

In this study, we demonstrate that standard-of-care therapies currently used in paediatric BRAF-V600E-mutant gliomas, including the MEK inhibitor Trametinib, the BRAF-V600E inhibitor Dabrafenib, and the chemotherapeutic agent Vinblastine, induce a senescence program in BT-40 tumour cells both in vitro and in vivo. Importantly, we show that this senescent state is associated with the activation of a SASP, leading to increased secretion of pro-inflammatory cytokines and recruitment of IBA1^+^ microglia within the tumour microenvironment. Notably, despite *CDKN2A/B* deletion in BT-40 cells, which prevents p16^INK4a^-dependent senescence ^12^, these tumour cells are able to activate a treatment-induced senescence program, most likely through CDKN1A (p21^Cip1^) expression, suggesting that MAPK pathway inhibition triggers a p53/p21-driven arrest in this model. We identify Navitoclax as a potent senolytic agent capable of eliminating therapy-induce senescent BT-40 cells and reducing or even preventing tumour rebound growth upon Trametinib withdrawal. Combination therapy with Trametinib and Navitoclax significantly reduces tumour burden and prolongs survival in a preclinical BT-40 orthotopic mouse model, highlighting a promising therapeutic strategy for paediatric gliomas carrying BRAF-V600E mutations. Our findings are in line with recent preclinical evidence showing that ERK inhibition with ulixertinib exerts potent antitumor activity in BRAF-driven pLGG models, including both proliferative and senescent states, supporting the idea that MAPK pathway blockade can efficiently modulate tumour cell fate independently of their baseline growth status ^38^.

Although all three drugs induced therapy-induced senescence (TIS), our transcriptomic analyses revealed drug-specific SASP profiles. Trametinib treatment induced stronger upregulation of lysosomal genes, extracellular matrix components (COL1A1, COL3A1, COL5A2), and cytokines such as CXCL6, CSF1, and TIMP2, whereas Dabrafenib and Vinblastine preferentially upregulated IL1A, IL1B, TGFB1, and MMP17. These observations are consistent with emerging evidence that TIS does not represent a uniform biological program but instead produces highly context-dependent SASP signatures, strongly influenced by the nature of the inducing agent^39^. For example, CDK4/6 inhibition can drive a SASP enriched in immune-stimulatory chemokines such as CXCL9, CXCL10, and MICB, enhancing NK and T cell recruitment ^40^, whereas genotoxic agents like docetaxel induce a pro-metastatic and immunosuppressive SASP ^41,42^. In our model, the distinct SASP profiles triggered by Trametinib, Dabrafenib, and Vinblastine likely contribute to differential immune modulation within the tumour microenvironment and may explain variability in therapeutic efficacy. Beyond TIS, the SASP is increasingly recognised as an intrinsic feature of pLGG biology driven by oncogene-induced senescence (OIS). Recent studies have shown that SASP signalling is already active in pLGG and may contribute to both tumour maintenance and microenvironmental crosstalk ^27^. This raises the possibility that MAPK inhibitor–induced senescence occurs within a cellular context already conditioned by OIS-associated SASP activity. In such a landscape, pharmacologically induced senescence may not only reinforce pre-existing SASP programmes but also amplify inflammatory and immune-modulatory signalling, potentially enhancing the microglial responses and tumour–stroma interactions we observe.

A striking finding of our study is that Trametinib-induced senescence was accompanied by the secretion of microglia-attracting chemokines and cytokines, including CCL2, CXCL10, and IL6, which correlated with an expansion of IBA1+ microglia within the tumour microenvironment. Although microglia have context-dependent roles in glioma progression, recent studies suggest that their recruitment by senescent cells may promote an immunosuppressive environment and facilitate tumour regrowth after treatment withdrawal ^19,43,44^. Our data showing persistent microglial infiltration even after Trametinib discontinuation, despite partial reduction of SASP cytokines, suggest that therapy-induced changes in the tumour microenvironment may predispose to tumour rebound. These findings are consistent with reports in paediatric low-grade gliomas demonstrating that MAPK pathway inhibition leads to compensatory immune responses, which may undermine long-term therapeutic efficacy^17,45^.

We identify Navitoclax, a well-characterised senolytic targeting BCL-2 family proteins, as a potent inducer of apoptosis in TIS BT-40 cells. Using selective inhibitors against individual anti-apoptotic proteins, we reveal that blockade of BCL-xL is the principal mechanism underlying Navitoclax’s activity. This aligns with previous reports showing that senescent cells depend on anti-apoptotic signalling for survival and are therefore preferentially susceptible to BCL-xL inhibition ^46^. Notably, treatment-induced senescent BT-40 cells displayed markedly lower IC50 values for Navitoclax than proliferative BT-40 cells, with Trametinib-induced senescent cells showing the greatest sensitivity. These findings parallel those of Sigaud et al., who reported strong synergy between the ERK inhibitor ulixertinib and BH3-mimetics, particularly the BCL-xL inhibitor Navitoclax, across multiple BRAF-driven pLGG models ^38^. Moreover, BCL-xL inhibition has previously been proposed as a therapeutic vulnerability in pLGG in the setting of OIS ^26^. Our results extend this vulnerability to TIS, suggesting that combining BCL-xL inhibition with MAPK pathway blockade may offer a dual strategy to eliminate both OIS- and TIS-associated senescent cell populations within the tumour.

In vivo, a combination therapy with Trametinib and Navitoclax leads to significant tumour regression, prolonged survival, reduced SASP cytokine expression, and decreased microglial infiltration, providing proof-of-concept for a senolytic-based therapeutic approach in paediatric gliomas. Our data also suggest that the benefits of Navitoclax are not merely due to tumour debulking but rather to the selective removal of senescent cells, thereby reducing pro-tumorigenic SASP-driven microenvironmental changes.

Our findings support a novel therapeutic paradigm in paediatric BRAF-V600E gliomas: combining MAPK pathway inhibitors with senolytics to enhance treatment efficacy and reduce tumour regrowth after drug withdrawal. This strategy may have relevance in preventing long-term dependence on targeted therapies, which can be associated with cumulative toxicity, quality-of-life impairment, and risk of malignant progression ^47^. However, translating our results into the clinic requires addressing several key challenges including toxicity concerns. Navitoclax is indeed associated with dose-limiting thrombocytopenia in paediatric and adult trials ^48^, developing BCL-xL-selective inhibitors or intermittent dosing schedules may improve tolerability^46^. Another aspect is patient heterogeneity, given the context-dependent nature of SASP profiles, identifying predictive biomarkers of both senescence induction and senolytic susceptibility will be essential to design therapeutic strategies. Moreover, blood-brain barrier (BBB) penetration of Navitoclax may be limited, although recent evidence in primates has demonstrated that this compounds can cross the BBB and accumulate in the cerebrospinal fluid at concentrations able to ablate senescent cells ^49^. Finally, this emerging ‘one-two punch’ therapeutic paradigm requires an optimum combination timing, determining the best therapeutic window for senolytic administration relative to MAPK inhibition remains an open question. Future studies should evaluate this combination strategy across diverse BRAF-mutant glioma models, integrate immune profiling to better understand microglial dynamics, and explore senolytic alternatives with improved safety profiles.

While our findings provide strong preclinical evidence, several limitations should be considered, including that our analyses are based primarily on the BT-40 cell line, which carries both a BRAF-V600E mutation and CDKN2A/B deletion. Broader validation in additional paediatric glioma models is warranted. Although we demonstrate SASP-driven microglial recruitment, the functional role of microglia, whether pro-tumorigenic or anti-tumorigenic, remains to be elucidated.

## Conclusion

Our study uncovers a previously unappreciated role of therapy-induced senescence in shaping the response of paediatric BRAF-V600E gliomas to current standard treatments. We demonstrate that senescent BT-40 tumour cells exhibit a selective vulnerability to BCL-xL inhibition, and that combining Trametinib with Navitoclax effectively reduces tumour burden, suppresses SASP-driven microenvironmental changes, and reduce or prevent tumour regrowth after Trametinib withdrawal. These findings provide a strong preclinical rationale for integrating senolytics into therapeutic regimens for paediatric gliomas carrying BRAF-V600E mutations.

## Authorship

RG and HH carried out most of the experiments and prepared all the figure panels for publication. AV, LC provided expertise and carried out brain cell injection. RS carried out the analysis shown in Figure 4D. SH, AB, DG, TS contributed experimentally. RC, JB, ML performed tumour imaging/irradiation. TJ, DH, JG, OW, TM provided expertise and intellectual input. The project was designed by RG and JPMB. RG and JPMB wrote the paper, with help from HH. All authors reviewed and approved this submission.

## Data availability

Raw and processed RNA sequencing data are available on GEO repository GSE294664 and GSE296000.

## Acknowledgements

We thank core facilities such as UCL Genomics, Light Microscopy facility and UCL Biological Services Unit. We thank Prof. Owen Williams (University College London) for providing LucF-GFP lentiviral vector. We thank our funders and sponsors for enabling the research presented in this manuscript.

## Supplementary data (separate file)

### Supplementary file

- Supplementary material
- Supplementary Figures S1-S6
- Supplementary Table S1-S2

Supplementary Table S3

Supplementary Table S4

Supplementary Table S5

Supplementary Table S6

Raw data

