## Supplementary material for "Standard treatment against paediatric BRAF-V600E glioma promotes senescence and sensitizes tumours to BCL-xL inhibition"

#### Supplementary Methods

##### BT-40 cell line culture

BT-40 cells (kind gift of Dr. P. Houghton, University of Texas) were grown in RPMI 1640 Medium, GlutaMAX™ Supplement (Gibco, catalogue no. 61870010) supplemented with 10% fetal bovine serum (Fetal Bovine Serum, qualified, Brazil, Gibco, catalogue no. 10270106, batch 2166437). Cells were routinely confirmed mycoplasma negative using the MycoAlert™ PLUS Mycoplasma Detection Kit (Lonza, catalogue no. LT07-710) and STR profiled.

##### Drugs for in vitro and in vivo testing

Dabrafenib (MedChemExpress [MCE] - HY-14660), Trametinib (ApexBio - A3018), Vinblastine (Santa Cruz Biotechnology - sc-491749), Navitoclax (ABT-262; ApexBio - A3007), A-1155463 (MCE - HY-19725), A-1331852 (MCE - HY-19741), Obatoclax (MCE - HY-10969), Venetoclax (ABT-199; ApexBio - A8194), Piperlongumine (ApexBio - A4455), Digoxin (ApexBio - B7684).

For in vitro use, drugs were dissolved in DMSO (Santa Cruz Biotechnology - sc-202581) and stored in aliquots at −20°C. Drugs were diluted in cell culture medium and added to the cells for the durations and concentrations indicated.

For in vivo experiment, Navitoclax powder was weighed, individual aliquots were prepared. Prior to administration, vehicle (ethanol:polyethylene glycol 400:Phosal 50 PG) was added and drug was sonicated on ice into solution using Diagenode Bioruptor®, high frequency, 30 second intervals, for 15 minutes. Navitoclax was administered to mice by gavage at 50 mg per kg body weight per day (mg/kg/d) for 5 days for two cycles. Trametinib was dissolved in a DMSO 20mg/mL stock solution, individual aliquots were prepared and stored at -20°C. Prior to administration, Trametinib stock solution was diluted 1:100 in vehicle (PBS 10%w/v (2-Hydroxypropyl)-β-cyclodextrine – Sigma Aldrich C0926). Trametinib was administered to mice by I.P. injections at 1 mg/kg/d.

**Generation of EGFP-LucF-BT-40 cells**

For viral particles production, 293FT cell line (Invitrogen) were transfected with the plasmids pSLIEW, pMD2.G and psPAX2 (Addgene plasmids 12259 and 12260) using Lipofectamine™ 2000 Transfection Reagent (Invitrogen). For cell transduction, BT-40 cells were seeded in 24 well plates, and transduced at 70% confluence. Medium was removed by aspiration and replaced with fresh complete medium containing 5 µg/ml polybrene (Millipore). Concentrated viral supernatant was added to achieve a multiplicity of infection (MOI) of 10. After 18 hours, the medium was replaced with fresh complete medium to remove lentivirus and polybrene. Transduced cells were expanded and sorted using flow cytometry assisted cell sorting (FACS), based on EGFP fluorescence. Sorted cells were expanded and stored in liquid nitrogen.

**Intracranial cell injections**

A single-cell suspension (EGFP-LucF-BT-40) was prepared immediately before implantation in 6 weeks old NOD-SCID-gamma female mice (Charles River UK). For intracranial procedures, animals were anesthetised with isoflurane. The cranium was exposed via midline incision under aseptic conditions and 1×1mm deep hole was made through the skull to the dura. Mice were placed in a stereotactic apparatus and 60,000 cells in 2 µL were stereotactically implanted into the right striatum area using a 10µL 26G Hamilton syringe at a rate of ~ 1 µL/min. Co-ordinates used were 1.5mm lateral to midline, 1.5mm posterior to bregma, and 4 mm deep to cranial surface. A small pocket was created for the cells, which were injected 3 mm deep to the cranial surface. At the completion of infusion, the syringe needle was allowed to remain in place for a minimum of 2 minutes, then slowly manually withdrawn to minimize backflow of the injected cell suspension. The skin was closed with VetBond™ Adhesive. Mice were weighed daily for one week, and dosed for 48 hours with Metacam, at 5 mg/kg.

**bulkRNA sequencing on BT-40 xenografts**

Once tumours were established, the mice received either Vehicle or Trametinib treatment during one week. Tumour bearing brain were dissected into 1 mm3 pieces and dissociated in a solution of 0.5% w/v collagenase type 2 (Lorne Laboratories Ltd.), 0.1x Trypsin (Gibco), 50 µg/ml DNaseI (Worthington) and 2.5 µg/ml fungizone (Gibco) in Hank's Balanced Salt Solution (HBSS (Gibco) for 45 minutes at 37°C. Tissue fragments were further dissociated into single cells through mechanical trituration using a pipette and filtration through a 70 µm filter. Single cells were washed in HBSS and suspended in FACS media (PBS supplemented with 1% foetal calf serum (PAA), 25 mM HEPES and 10 µg/ml of 4′,6-diamidino-2-phenylindole (DAPI, BD Pharmingen) for live/dead cell discrimination) before being analysed using a BD FACS AriaIII Flow Cytometer. GFP+ cells were collected into 4°C RLT lysis buffer and total RNA was isolated (Qiagen RNeasy Micro Kit). Total RNA was then processed for bulk RNA-sequencing analysis. Isolated RNA was processed by the SMARTer V4 low input assay kit (Clontech) to generate amplified cDNA using the strand-switching protocol. Library preparation was performed with 200 pg of amplified cDNA using the NEBNext Low Input RNA kit with 12 cycles of PCR. Sequencing was then performed on an Illumina NextSeq 2000 100cycles with 100 bp single-end reads. To remove potential contamination of host cells from the analysis, sequencing data was aligned with {STAR 2.7.11b} on both human and mouse annotated genomes (Ensembl genome 113). {XenofilteR 1.6} package (Kluin RJC, Kemper K, Kuilman T, et al. XenofilteR: computational deconvolution of mouse and human reads in tumor xenograft sequence data. *BMC Bioinformatics*. 2018;19(1):366. doi:10.1186/s12859-018-2353-5) was used to filtered out murine host reads from human graft reads. Raw and processed data are available on GEO repository GSE296000. Secondary analysis was performed using {R 4.3.3} on RStudio (2023.06.0+421). Differential expression was performed using {DESeq2 1.38.1}, complete pipeline is available at https://gitlab.univ-nantes.fr/guihomics/plgg_tis_rnaseq. GSEA performed using the GSEA software (v4.3.2 - UC San Diego and Broad Institute). Gene sets used in this study were either obtained from public databases (gene ontology (GO), KEGG, REACTOME, molecular signatures database (Broad Institute) or published literature: senescence profiling and SASP.

**Assessment of apoptosis**

Two thousand cells per well were seeded into 96-well plates and cultured for 24 h in the presence of 100nM Vinblastine, Dabrafenib or Trametinib as indicated, followed by a wash out period of 6 days. At day 7, half of the plates were fixed with PFA 4%, stain with DAPI, scanned on Zeiss Observer 7 colour fluorescence and nuclei were counted with Fiji software 1.53. On the other half, Navitoclax (0, 0.1 or 1µM) was added to the pre-treated cells for 24h. Caspase 3/7 activity was determined using 100µl Caspase-Glo® 3/7 Assay System (Promega G8090), which was added directly to the cells in 96-well plates and incubated for 1 hour before recording luminescence on a Microplate luminometer (GloMax® Navigator – Promega). The "no cell" blank control value has been subtracted from each, and results were normalized by DAPI-positive nuclei counted with the same treatments conditions.

### Supplementary Figures

#### Suppl. Figure S1

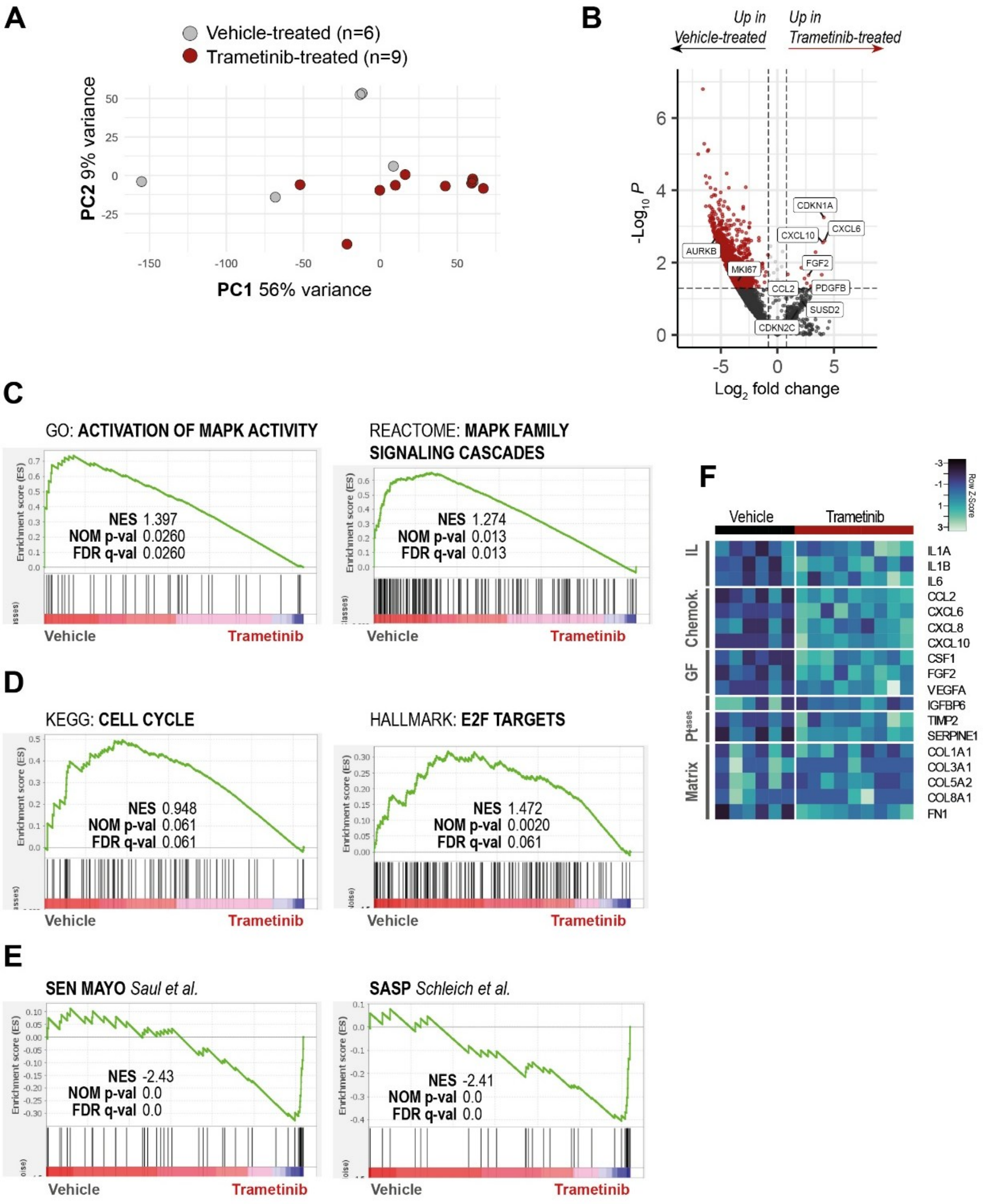

**Supplementary Figure S1. Transcriptomic analysis of BT-40 orthotopic xenograft tumours treated with the MAPK inhibitor Trametinib confirms MAPK pathway inhibition and cell cycle arrest, while revealing the induction of senescence-associated features.** (A) Principal component analysis plot of the 6 Vehicle-treated (grey) and the 9 Trametinib-treated (red) samples based on single-end RNAseq of flow-sorted BT-40 tumour cells. (B) Volcano plot of differentially expressed genes identified between vehicle and Trametinib-treated group. Thresholds used: p-values < 0.05 and log2 FoldChange > 1. (C-E) Geneset enrichment analysis suggest the inactivation of MAPK pathway (C), the decrease in proliferative function (D) and the enrichment for senescence and senescence-associated secretory phenotype (E) in Trametinib-treated tumour cells. (F) Heatmap highlighting the diversity of genes coding for secretory and matrix proteins associated with SASP overexpressed in the Trametinib-treated samples.

#### Suppl. Figure S2

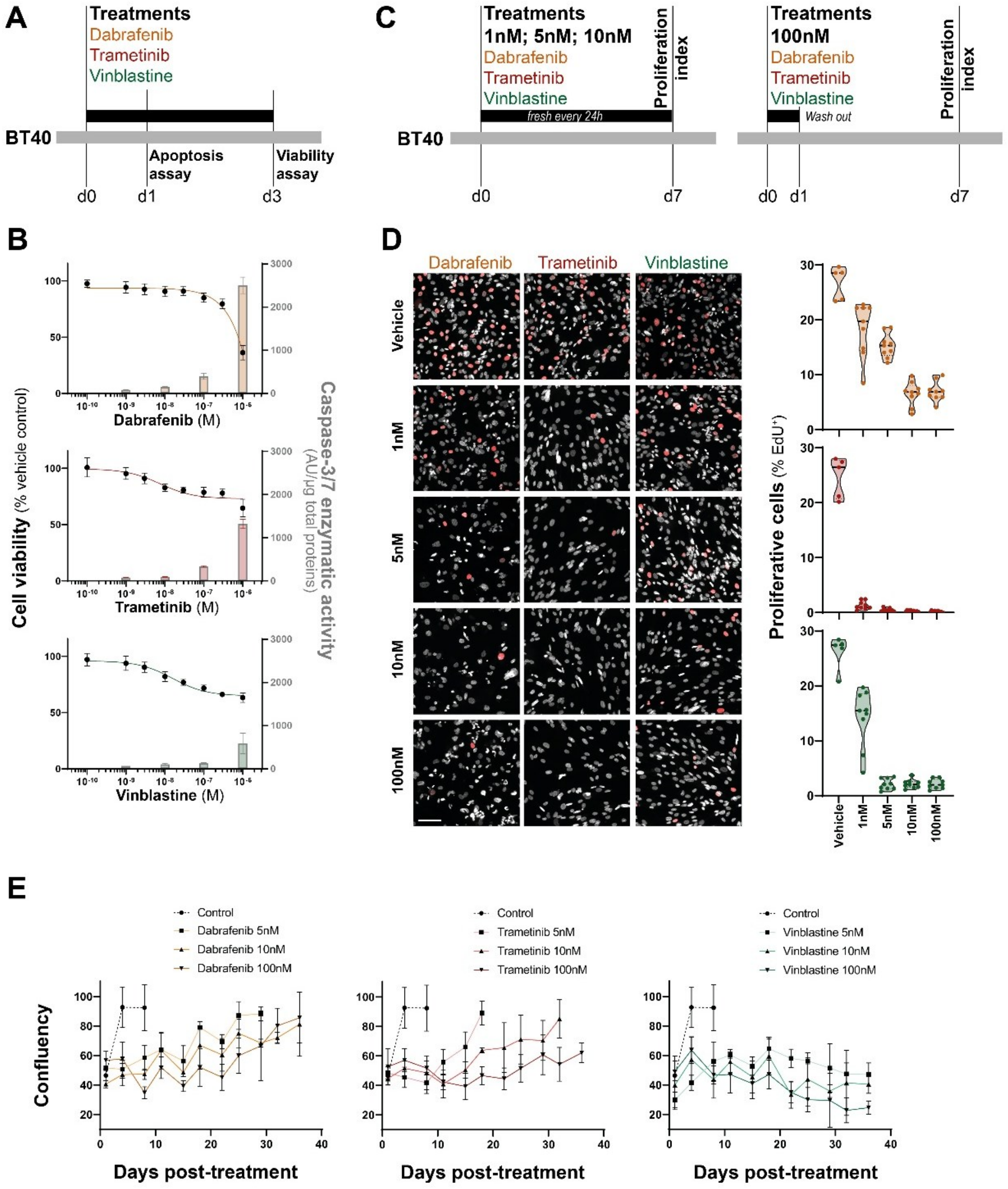

**Supplementary Figure S2. In vitro treatment with MAPK inhibitors or Vinblastine induces sustained proliferation arrest in BT-40 tumour cells.**

(A) A schematic of treatments and analysis kinetic for short term effect of a range of concentrations. (B) BT-40 cell viability assessed by ATP production at day 3 (line) and apoptosis assessed by caspase 3/8 activity at day 1 (bar) in the presence of increasing concentrations of Trametinib, Dabrafenib or Vinblastine. (C) A schematic of treatments and analysis kinetic for long term effect a chronic treatment of 1, 5 or 10nM of the drugs or a 24h pulse of 100nM on BT-40 proliferation index. (D) EDU accumulation overnight observed by Click-iT kit indicating proliferation of the treated cells at Day 7. Quantification of % of positive nuclei on the right. (E) Assessment of confluency overtime showing an overcoming of cell proliferation arrest 2 to 3 weeks post MAPK inhibitors treatment whereas vinblastine-treated cells remain non-proliferative for more than 5 weeks. Shown are mean ± SEM of three independent experiments.

#### Suppl. Figure S3

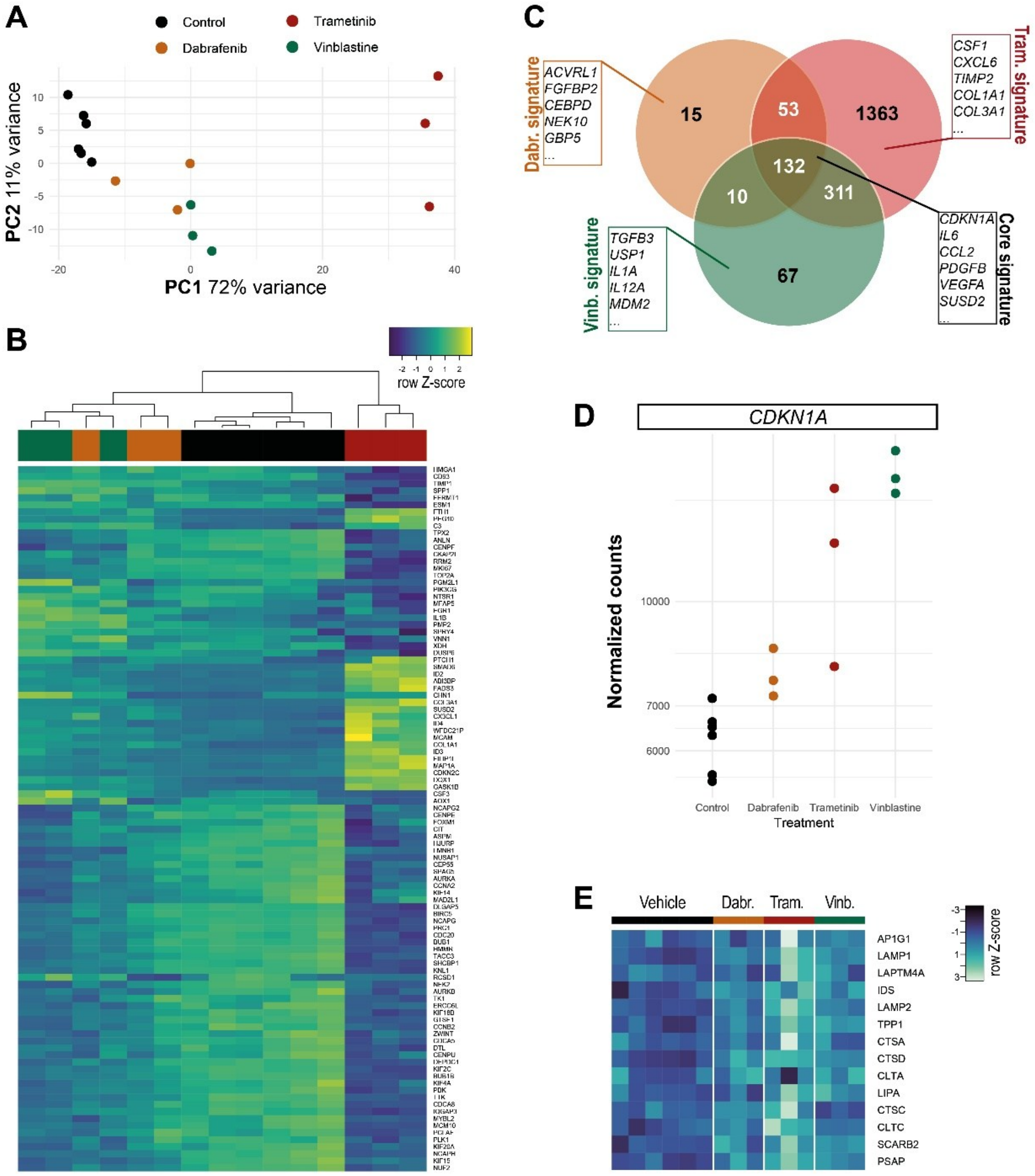

**Supplementary Figure S3. Transcriptomic analysis by RNAseq of Treatment-induced senescent BT-40 cells reveals both shared and drug-specific gene expression signatures.**

(A) Principal component analysis plot of the 6 proliferative controls (black), the 3 Trametinib-induced senescent (red), the 3 Dabrafenib-induced senescent (orange) and the 3 Vinblastine-induced senescent (green) samples based on RNAseq of BT-40 tumour cells. (B) Heatmap of the 500 most variable genes enabling segregation between the different conditions. (C) Venn-diagram of the number of differentially expressed genes identified between proliferative and treatment-induce senescent state, classified according to inducing treatment, revealing core and treatment-specific gene signatures. (D) Difference in *CDKN1A* ($p21^{CIP1}$ coding gene) normalized counts across the proliferative and treatment-induced senescence states. (E) Heatmap highlighting the diversity of genes coding for proteins associated with lysosomal compartment overexpressed in the treatment-induced senescence samples.

**Suppl. Figure S4**

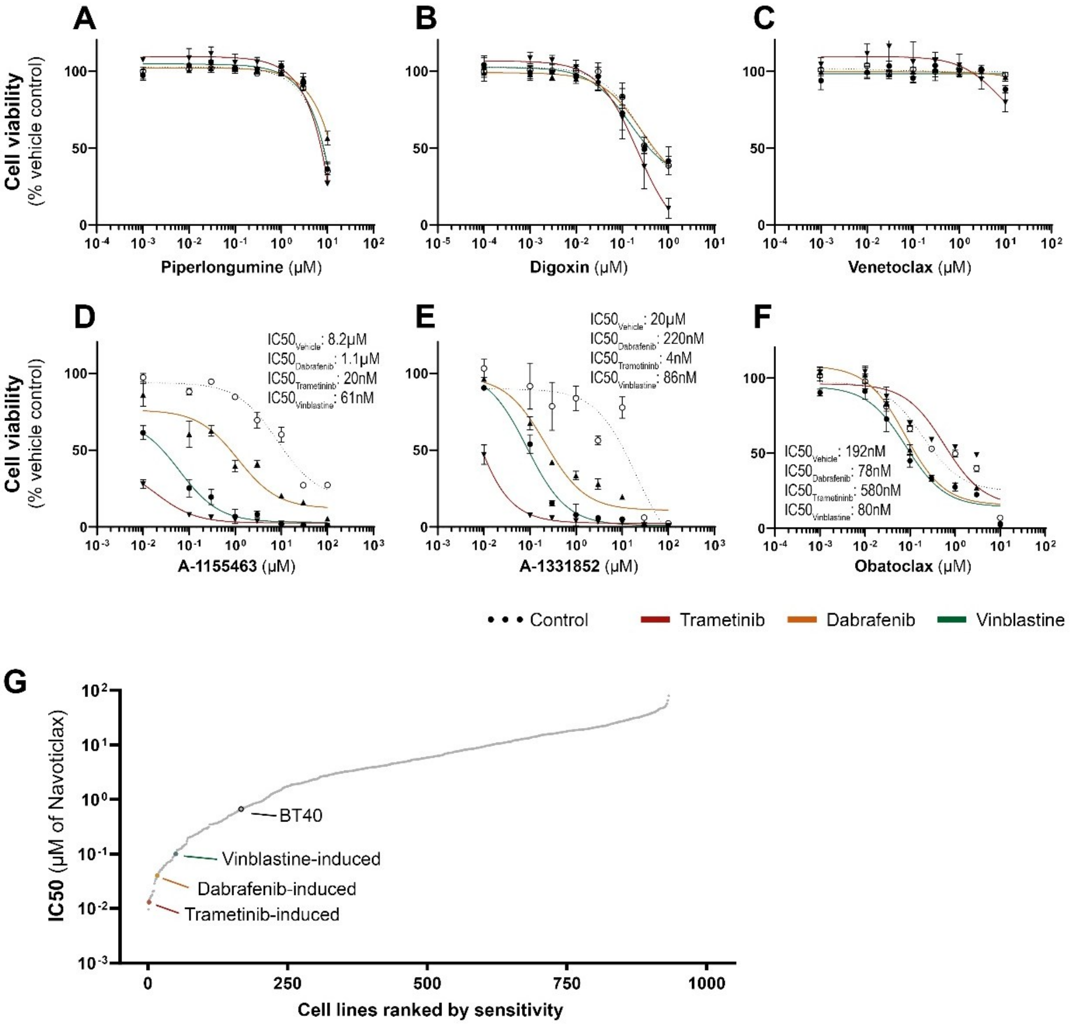


**Supplementary Figure S4. Treatment-induced senescent BT-40 cells are selectively sensitive to the Bcl-xL inhibitors A-1331852 and A-1155463.** Assessment of viability by ATP-production of treatment-induced senescent (Dabrafenib: orange, Trametinib: red, Vinblastine: green) or proliferating (dotted black) BT-40 cells treated for 72 hours with known senolytics (Piperlongumine (A), Digoxin (B)) or BH3-mimetics Venetoclax (C), A-1155463 (D), A-1331852 (E) and Obatoclax (F) at indicated concentrations. Shown are mean ± SEM of three independent experiments. 928 cell lines screened by the Genomics of Drug Sensitivity in Cancer (GDSC) cell lines database ranked by Navitoclax IC50. Parental BT-40, not included in GDSC database, and treatment-induced senescent BT-40 cells have been inserted into their corresponding positions in the ranking according to experimentally calculated IC50: 2nd for trametinib-induced senescence, 17th for dabrafenib-induced senescence, and 50th for vinblastine-induced senescence (G).

#### Suppl. Figure S5

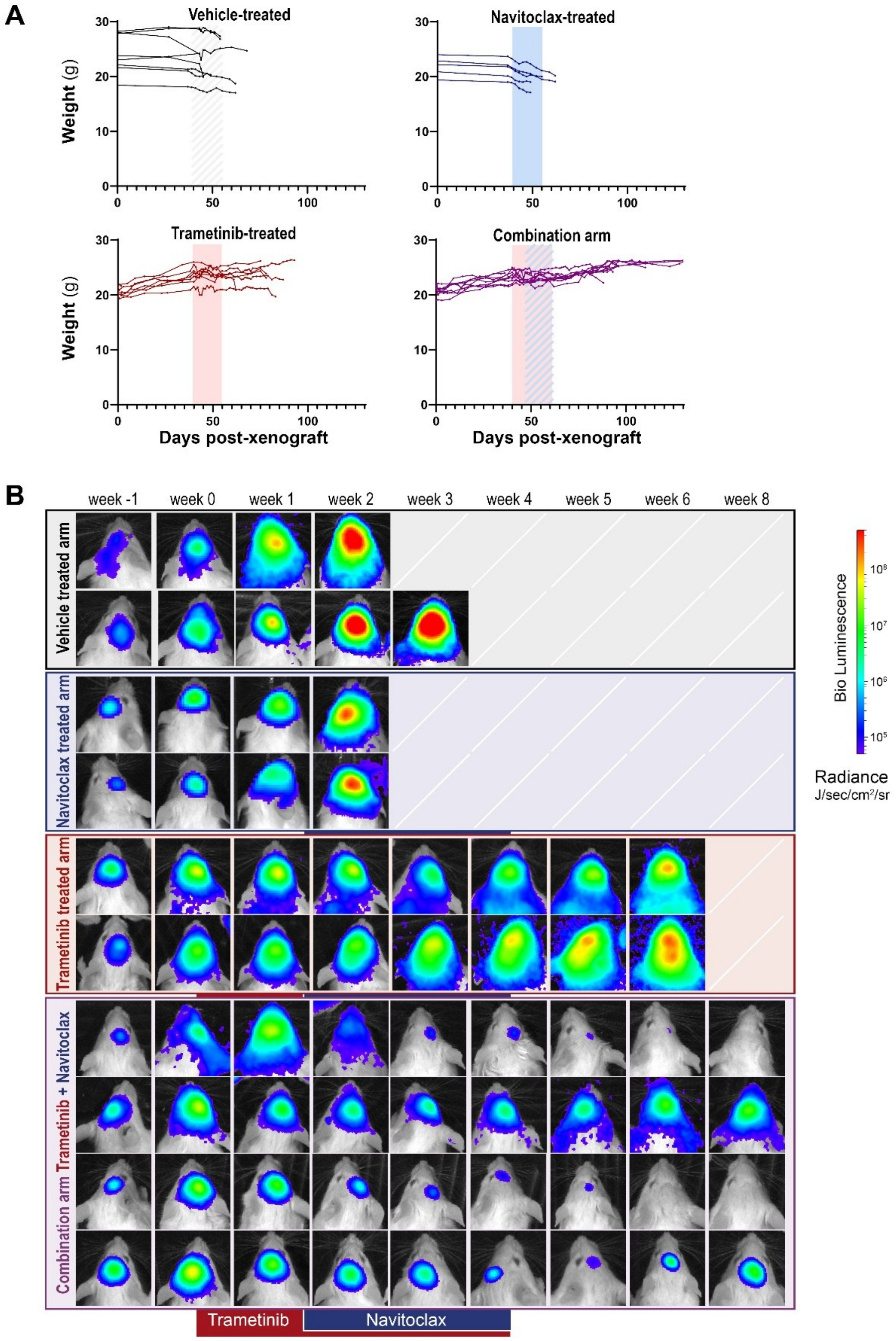

**Supplementary Figure S5. Individual follow-up of mice bearing orthotopic BT-40 xenograft tumours treated with Trametinib, Navitoclax, or sequential Trametinib followed by combination therapy.** (A) Weight of each animal per group. Human endpoint is fixed when weight drop by >10% ; (B) BLI representation across the time, before, during and after treatments for representative animals in each group.

#### Supplementary Figure S6

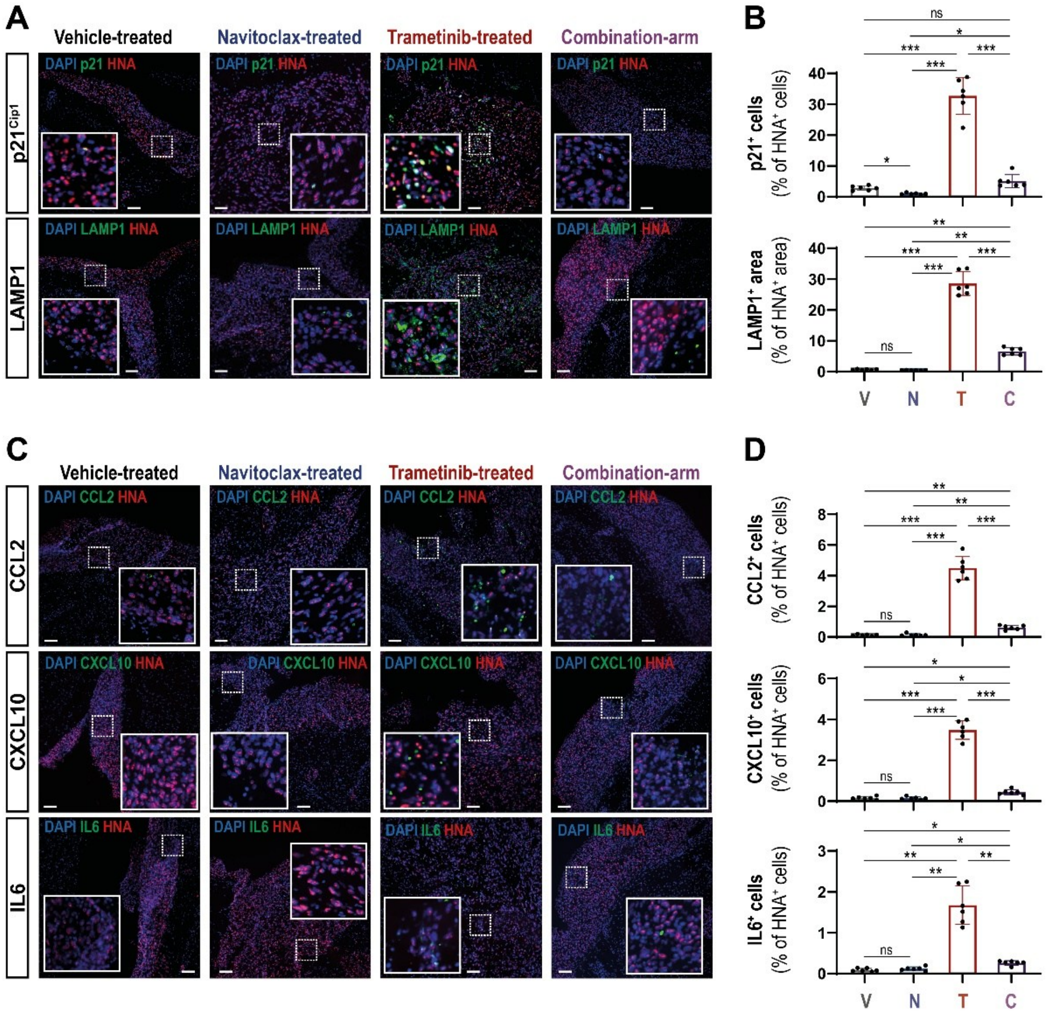


**Supplementary Figure S6. Combination therapy with Trametinib and Navitoclax reduces senescence and SASP marker expression in BT-40 tumour cells in vivo.**

(A) Double immunofluorescence against Human Nuclear Antigen (HNA –red) and the senescence markers p21 or LAMP1 (green). Scale bars: 200µm. (B) Quantification of the proportion of HNA positive cells expressing p21 and LAMP1 positive tumour area analysed in (A). Note the overall reduction of senescent marker expression in the combination compared with the Trametinib-only arms. (C) Double immunofluorescence against Human Nuclear Antigen (HNA –red) and the SASP markers CCL2, CXCL10 or IL6 (green). Scale bars: 200µm. (D) Quantification of the proportion of HNA positive cells expressing the markers used in (C). Note the overall reduction of SASP marker expression in the combination arm compared to Trametinib-only treated arm. Data show mean ± SEM of n = 3 BT-40 tumours per marker. Kruskal-Wallis test, Dunn's post-test; *: $p<0.05$, **: $p<0.01$, ***: $p<0.001$

**Supplementary Table S1. BH3 mimetics used in this study.**

| Compound | Inhibitory profile ($K_i$ cell free assays) | | | | Clinical trial status | Example clinical trial | Achievable non-toxic $C_{max}$ (nM) |
|---|---|---|---|---|---|---|---|
| | Bcl-2 | Bcl-xL | Bcl-w | Mcl-1 | | | |
| **Venetoclax** | <0.01 | 48 | 245 | >440 | MHRA approved | NCT04401748 | 2533.4 ± 1612 |
| **Navitoclax** | <1 | <0.05 | <1 | 550 (+/-40) | Phase III | NCT02591095 | 6607 ± 3262 |
| **Obatoclax** | 0.2 | 1-7 | 1-7 | 1-7 | Phase I | NCT00600964 | 7-38 |
| **A-1331852** | 6 | <0.01 | 4 | 142 | Preclinical | n.a | n.a. |
| **A-1155463** | 74 | <0.01 | 8 | >444 | Preclinical | n.a | n.a |

**Supplementary Table S2. Primary and secondary antibodies used in this study.**

| Application | Target | Host | Clonality | Clone | Supplier | Catalogue No | Dilution | Antigen Retrieval Method | Conjugate |
|---|---|---|---|---|---|---|---|---|---|
| **Primary** FFPE | Human nuclei antigen | Mouse | Monoclonal | 3E1.3 | Sigma-Aldrich | MAB4383 | 1:100 | Tris- EDTA pH9.0 | |
| **Primary** FFPE | Ki67 | Rabbit | Monoclonal | SP6 | Abcam | ab16667 | 1: 500 | Tris-EDTA pH9.0 | |
| **Primary** In vitro | p21 | Rabbit | Polyclonal | M-19 | Cell Signaling Technology | #5487 | 1:400 | Tris- EDTA pH9.0 | |
| **Primary** FFPE | p21 | Rabbit | Polyclonal | N/A | eBioscience | 14-6715-81 | 1:100 | Tris-EDTA pH9.0 | |
| **Primary** FFPE | p21 | Mouse | Monoclonal | SXM30 | BD Pharmingen | 6431 | 1:100 | Tris-EDTA pH9.0 | |
| **Primary** FFPE | Lyzozomal β-Galactosidase (GLB1) | Rabbit | Polyclonal | N/A | Proteintech | 15518-1AP | 1:100 | Tris-EDTA pH9.0 | |
| **Primary** FFPE | LAMP-1 | Rabbit | Polyclonal | N/A | Abcam | ab42170 | 1:200 | Tris-EDTA pH9.0 | |
| **Primary** In vitro | CCL2 | Rabbit | Monoclonal | N/A | R&D | MAB679R | 1:1000 | Tris- EDTA pH9.0 | |
| **Primary** FFPE | CCL2 | Rabbit | Polyclonal | N/A | Invitrogen™ | 15365431 | 1:100 | Tris-EDTA pH9.0 | |
| **Primary** In vitro | IL1B | Rabbit | Monoclonal | N/A | R&D | MAB201 | 1:200 | Tris- EDTA pH9.0 | |
| **Primary** FFPE | CXCL10 | Rabbit | Polyclonal | N/A | Proteintech | 10937-1-AP-20UL | 1:100 | Tris-EDTA pH9.0 | |
| **Primary** FFPE | IL6 | Goat | Polyclonal | N/A | R&D systems | AF-406-N | 1:100 | Tris-EDTA pH9.0 | |
| **Primary** FFPE | IBA1 | Rat | Monoclonal | EPR16589 | Abcam | ab283346 | 1:100 | Tris-EDTA pH9.0 | |
| **Primary** FFPE | CD68 | Rabbit | Polyclonal | N/A | Aviva Systems Bio | OABB00472 | 1: 500 | Tris-EDTA pH9.0 | |
| **Primary** FFPE | Lamin B1 | Rabbit | Polyclonal | N/A | Abcam | ab16048 | 1: 500 | Tris-EDTA pH9.0 | |
| **Primary** FFPE | Cleaved Caspase | Rabbit | Polyclonal | Asp175 | Cell Signaling | #9661 | 1:1000 | Tris- EDTA pH9.0 | |
| **Secondary** | Mouse Ig | Goat | Polyclonal | Thermo Fisher | A11001 | | 1:250 | | Alexa Fluor® 488 |
| **Secondary** | Mouse Ig | Goat | Polyclonal | Thermo Fisher | A11004 | | 1:250 | | Alexa Fluor® 568 |
| **Secondary** | Rabbit Ig | Goat | Polyclonal | Thermo Fisher | A11008 | | 1:250 | | Alexa Fluor® 488 |
| **Secondary** | Rabbit Ig | Goat | Polyclonal | Thermo Fisher | A11036 | | 1:250 | | Alexa Fluor® 568 |
